# A microRNA CRISPR screen reveals miR-483-3p as an apoptotic regulator in prostate cancer

**DOI:** 10.1101/2025.04.16.649147

**Authors:** Jonathan Tak-Sum Chow, Ayeisha Desjardins, Daniel K.C. Lee, Iulia A Grigore, Norman J Fu, Stephanie Chau, Byeong Yeop Lee, Martino Marco Gabra, Leonardo Salmena

## Abstract

The development of traditional protein-targeted cancer therapies is a slow and arduous process, often taking years or even decades. In contrast, RNA-based therapies targeting crucial microRNAs (miRNAs) offer a faster alternative due to the sequence specific nature of miRNA inhibitor binding. This, combined with the capacity of individual miRNAs to influence multiple cellular pathways, makes these small RNAs attractive targets for cancer therapy. While miRNA are known to be dysregulated in prostate cancer (PCa), identifying their individual contributions to disease progression and the identification of therapeutically actionable miRNA targets in PCa has been challenging due to limited screening tools. To overcome this, we developed miRKOv2, a miRNA-only CRISPR knockout library enabling systematic, genome-wide loss-of-function screens to identify miRNAs essential for PCa cell survival. Our screens uncovered 69 potential essential miRNA candidates, with miR-483 demonstrating the most significant impact on PCa cell viability. Functional characterization demonstrated that miR-483 disruption significantly potentiated apoptosis in PCa cell lines. Mechanistically, we uncovered a novel regulatory axis wherein miR-483-3p directly modulates a BCLAF1/PUMA/BAK1 apoptotic signaling network, highlighting its critical role in maintaining PCa cell survival. Our findings provide novel insights into the complex regulatory role of miRNA in PCa progression and offer a potential therapeutic strategy for targeting miRNA-mediated pathways in metastatic disease.

## INTRODUCTION

Prostate cancer (PCa) is a prevalent solid malignancy among men around the world, resulting in over 1.5 million new diagnoses, and nearly 400,000 deaths each year (1,2). The treatment paradigm for metastatic PCa has greatly improved over the last decade, owing to an improved understanding of androgen receptor signaling, and the development of targeted androgen therapies (2–4). Notwithstanding the advancements in therapeutic modalities that have yielded improvements in survival outcomes, metastatic PCa remains an intractable and fatal disease, thereby underscoring a pressing requirement for enhanced therapeutic interventions.

MicroRNA (miRNA), typically 19 to 25 nucleotides long, are small endogenous RNAs that regulate gene expression post-transcriptionally. miRNA are thought to be crucial regulators of cellular homeostasis because they can control an estimated 60% of all protein-coding genes (5), and therefore almost every cellular process (6). As a result, dysregulation of miRNA is associated with many disease states, including cancer (7,8). Importantly, the sequence-specific binding of miRNA inhibitors enables precise targeting, making miRNAs viable drug targets. Furthermore, exogenous administration of miRNAs can compensate for loss or mutation in disease, collectively positioning them as attractive therapeutic options for cancer (9).

PCa often exhibits dysregulation of miRNA processing, characterized by altered levels of miRNA processing proteins DGCR8 and DICER (10–12). Additionally, individual miRNA can be disrupted, where gain or loss of expression have been implicated in promoting metastatic PCa. For instance, enhanced miR-194-mediated repression of SOCS2 has been shown to drive PCa metastasis, while inhibition of miR-194 suppresses PCa cell invasion (13). Treatment with inhibitors against miR-221 and miR-222 resulted in the derepression of p27 and reduced the growth of *in vivo* xenografts (14). Similarly, modified miR-21 inhibitors reduced tumour growth *in vivo* and remained stable in circulation (15). Together these examples underscore the feasibility and significant therapeutic potential of miRNA-based therapies in cancer.

Despite the established link between miRNA dysregulation and PCa progression, identifying their individual contributions to disease progression has been challenging due to limited screening tools. This has hindered the full realization of their potential as therapeutic targets. Consequently, there is a need for the development of tools to functionally investigate miRNA roles in cancer. To address this gap, we developed a miRNA-focused CRISPR library called the **<u>miR</u>**NA-only **<u>K</u>**nock**<u>O</u>**ut (miRKO) version-2 (miRKOv2), a second iteration of the library that builds upon the original miRKO library (16).

To identify novel essential miRNA in PCa cells, we employed miRKOv2 to screen DU145 and LNCaP cells. We identified 69 candidate essential miRNA hits and further validated miR-483 as a putative essential miRNA in several PCa cell models through its regulation of apoptosis. Specifically, we uncovered a novel regulatory axis wherein miR-483-3p directly regulates a BCLAF1/PUMA/BAK1 apoptotic signaling network. These findings provide evidence for miR-483-3p as a potential therapeutic target in PCa, particularly in the context of metastatic PCa, where apoptosis evasion is a significant clinical challenge (17,18).

## RESULTS

### Generation of miRKOv2, and enhanced miRNA-only CRISPR-KO library

To identify miRNA that are essential for PCa cell fitness we developed miRKOv2, a guide RNA library that incorporates the most updated on- and off-target gRNA scoring metrics (19) and a focus on targeting pre-miRNA hairpin stem regions to improve efficiency of miRNA disruption (20), and positive control gRNA targeting validated core essential and non-essential protein coding genes (21–23). We also included non-targeting gRNA as negative controls to make up 1% of the final library, to improve detection power. miRKOv2 consists of 6532 gRNA targeting 1649 pre-miRNA hairpins (3 or 4 gRNA per pre-miRNA), 800 gRNA targeting 100 core-essential and 100 non-essential genes (4 gRNA per gene), and 74 non-targeting gRNA. When benchmarked against other existing miRNA libraries (16,24,25) miRKOv2 exhibited comparable on-target and off-target profiles, however miRKOv2 strictly adheres to a 0.2 on-target score threshold, ensuring the exclusion of potentially inefficient gRNAs. This minimizes the risk of false negative results due to inefficient gRNA activity (Figure 1A). Notably, miRKOv2 has the fewest off-target protein-coding loci with a CFD off-target score > 0.2 which minimizes the likelihood of false positive results due to gRNA off-targets (Figure 1B).

**Figure 1.**
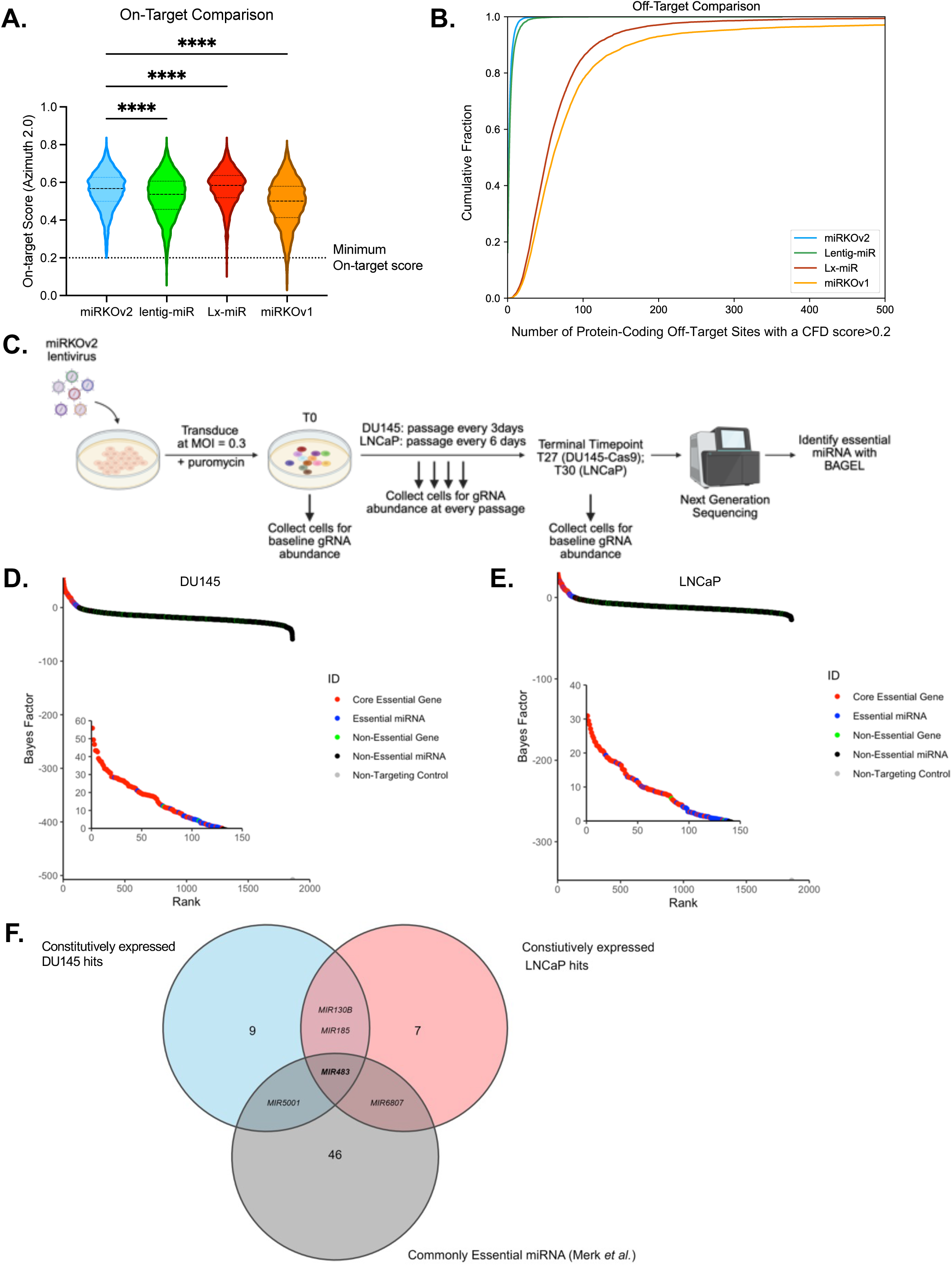
Identification of essential miRNA in PCa cells using a novel miRNA-focused CRISPR library. **A** On-target score comparison for gRNAs in the miRKOv2, lentiG-miR, Lx-miR and miRKOv1 libraries. Dotted line represents the recommended minimum on-target score of 0.2. Data represented by violin plots showing the first quartile, median and third quartile. **** *p* < 0.0001; One-way ANOVA with a Holm-Sidak’s multiple comparisons test. **B.** Comparison of cumulative off-target loci with a CFD score > 0.2 for the miRKOv2, lentiG-miR, Lx-miR and miRKOv1 libraries. **C.** Schematic representation of the miRKOv2 dropout screen in DU145-Cas9 and LNCaP cells. The screen was maintained at a 1000X gRNA representation and sequenced at a 500X gRNA representation. **D.** Genes ranked hits by Bayes Factor (BF) at the T27 terminal timepoint for the DU145-Cas9 dropout screen. Hits with a BF > 0 and FDR < 0.1 were considered essential. Essential hits are shown in the inset. **E.** Same as (**D**) but for the T30 terminal timepoint for the LNCaP dropout screen. **F.** Overlap of constitutively expressed hits from the DU145-Cas9 and LNCaP screens and commonly essential genes curated by Merk *et al.*(*24*) See also Figure S1 and File S1.

### Identification of essential miRNA in PCa cells

Dropout screens were performed in DU145 and LNCaP cells using miRKOv2 with day 27 and 30 days as the terminal time point, respectively (Figure 1C). Using BAGEL, a supervised learning algorithm that employs Bayesian statistics (26), we identified 32 essential miRNA hits in the DU145 screen and 47 essential miRNA in the LNCaP screen, with 10 common miRNA identified in both screens for a total of 69 unique hits at the terminal time point which performed the best in both screens (Figure 1D,E and S1A,B and File S1). Notably, many miRNA hits had BAGEL (Bayes Factor) scores that were comparable to core-essential coding gene controls, providing strong support for the notion that miRNAs can indeed function as crucial essential genes (Figure 1D,E insets).

For miRNA candidate prioritization, we cross-referenced miRNA expression profiling of a panel of PCa cell lines (DU145, PC3, 22Rv1, LNCaP and VCaP) with the screen candidate list.

We identified that 13/32 DU145 hits and 11/47 LNCaP hits were highly expressed (File S2). Furthermore, others have completed screens for essential miRNA, permitting the curation of a list of commonly essential miRNA across several cancer types (24). Integrating this list with our own list of screen candidates, we found *MIR483* to be commonly essential across multiple cancer types (24) (Figure 1F and File S2). Given the limited understanding of miR-483 function in PCa, our study aimed to validate and elucidate a potential mechanism responsible for its essentiality.

### MIR483 is essential for PCa cell growth

To validate *MIR483* as an essential gene, we assessed *MIR483*-targeting gRNAs in the miRKOv2 screens to reveal that three of the four gRNA were depleted at the terminal timepoints in both screens (Figure S2A,B). We also demonstrated that an independent *MIR483*-targeting gRNA distinct from those in miRKOv2 was also able to reduce cell growth in DU145, LNCaP, 22Rv1 and PC3 cells (Figure 2A-D). Together these data support that *MIR483* is an essential gene in PCa cells across diverse genetic backgrounds. Next, we observed that *MIR483* knockout in the premalignant immortalized prostate cell line RWPE-1 (27,28) did not result in any significant changes in cell growth (Figure 2E) suggesting that *MIR483* essentiality is restricted to malignant PCa cells.

**Figure 2.**
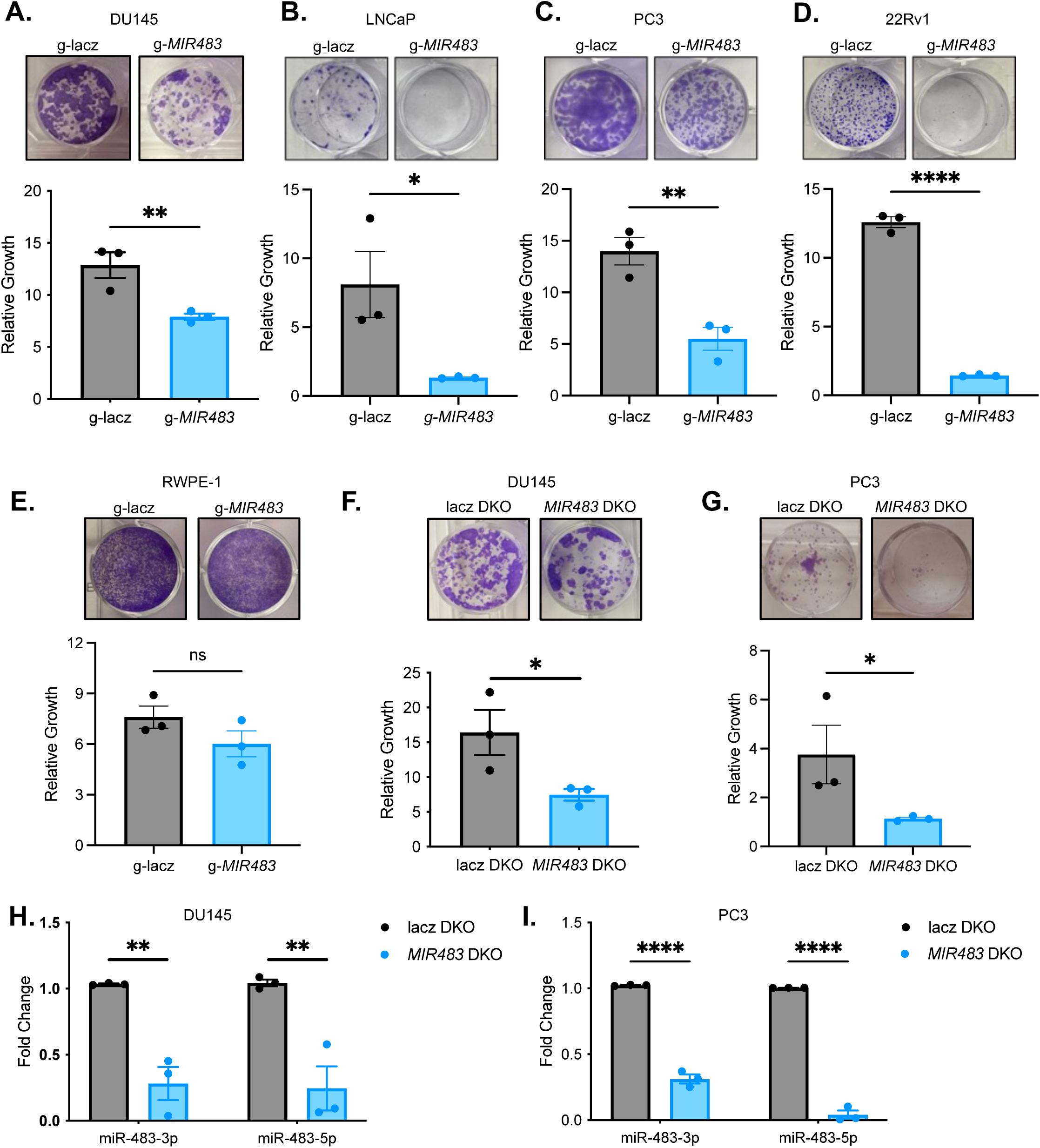
*MIR483* is essential for PCa cell growth. **A.** Growth assays of miRKOv2-independent gRNA targeting *MIR483* in DU145-Cas9 cells Representative images are shown on top and quantitation are shown on the bottom. **B.** Same as (**A**) in LNCaP cells. **C.** Same as (**A**) in PC3-Cas9 cells. **D.** Same as (**A**) in 22Rv1 cells. **E.** Same as (**A**) in RWPE-1 cells. **F.** Growth assay of *MIR483* DKO in DU145-Cas9 cells. Representative images are shown on top and quantitation are shown on the bottom. **G.** Same as (**F**) in PC3-Cas9 cells. **H.** RT-qPCR of miR-483-3p and miR-483-5p following *MIR483* knockout with the DKO system in DU145-Cas9 cells. **I.** Same as (**H**) in PC3-Cas9 cells. All data is represented as mean ± SEM from n=3 independent experiments. P-values obtained using an Unpaired one-tailed Student’s t-test (**A-F**) or a Two-way ANOVA with a Holm-Sidak’s multiple comparisons test (**H,I**). * *p* < 0.05; ** *p* < 0.01; **** *p* < 0.0001; ns, not significant. See also Figure S2 and File S2.

To independently validate these findings, a dual gRNA knockout (DKO) system was employed, revealing growth disruption in DU145 and PC3 cell lines and further confirming that *MIR483* is essential for prostate cancer (PCa) cell proliferation (Figures 2F, G). The DKO system was specifically designed to generate two indels flanking the *MIR483* genomic locus, thereby completely excising the miRNA sequence.(Figure S2C). Using RT-qPCR, we confirmed the DKO-mediated depletion of mature miR-483-3p and miR-483-5p (Figures 2H,I). Tracking of Indels by Deconvolution (TIDE) analysis (29) further revealed a 40-45 nucleotide deletion within the *MIR483* locus (Figure S2D,E), corresponding to the pre-miR-483 sequence, demonstrating successful DKO-mediated disruption.

Furthermore, recognizing that *MIR483* resides within the second intron of *IGF2*, a gene encoding a peptide hormone essential for growth (30,31), we ensured that *MIR483* knockout did not inadvertently disrupt *IGF2* expression. Indeed, IGF2 protein levels remained unchanged after *MIR483* knockout, ruling out *IGF2* disruption as a confounding factor in our experiments (Figure S2F,G). In sum, we identified and provided definitive evidence that *MIR483* is an essential gene in PCa cells through orthogonal approaches.

### miR-483 targets apoptotic pathways and its knockout increases apoptosis in PCa cells

The mechanisms underlying miR-483 essentiality were further explored using the Pathway Data Integration Portal (PathDIP) and Gene set enrichment analysis (GSEA). PathDIP revealed an enrichment of apoptosis-related pathways downstream of miR-483 (Figure 3A). GSEA was performed on publicly available RNA-sequencing data from control and *MIR483* knockout cell lines to further investigate the gene expression changes associated with miR-483 alterations in diverse cell lines (24). This analysis revealed that pathways associated with intrinsic apoptosis -- as identified in the KEGG (32) and ACSN2 (33) databases -- were notably enriched in *MIR483* knockout HT29, PC9, and MCF7 cells (Figure S4). Interestingly, pathways associated with normal mitochondrial processes were enriched in control cells (Figure S4).

**Figure 3.**
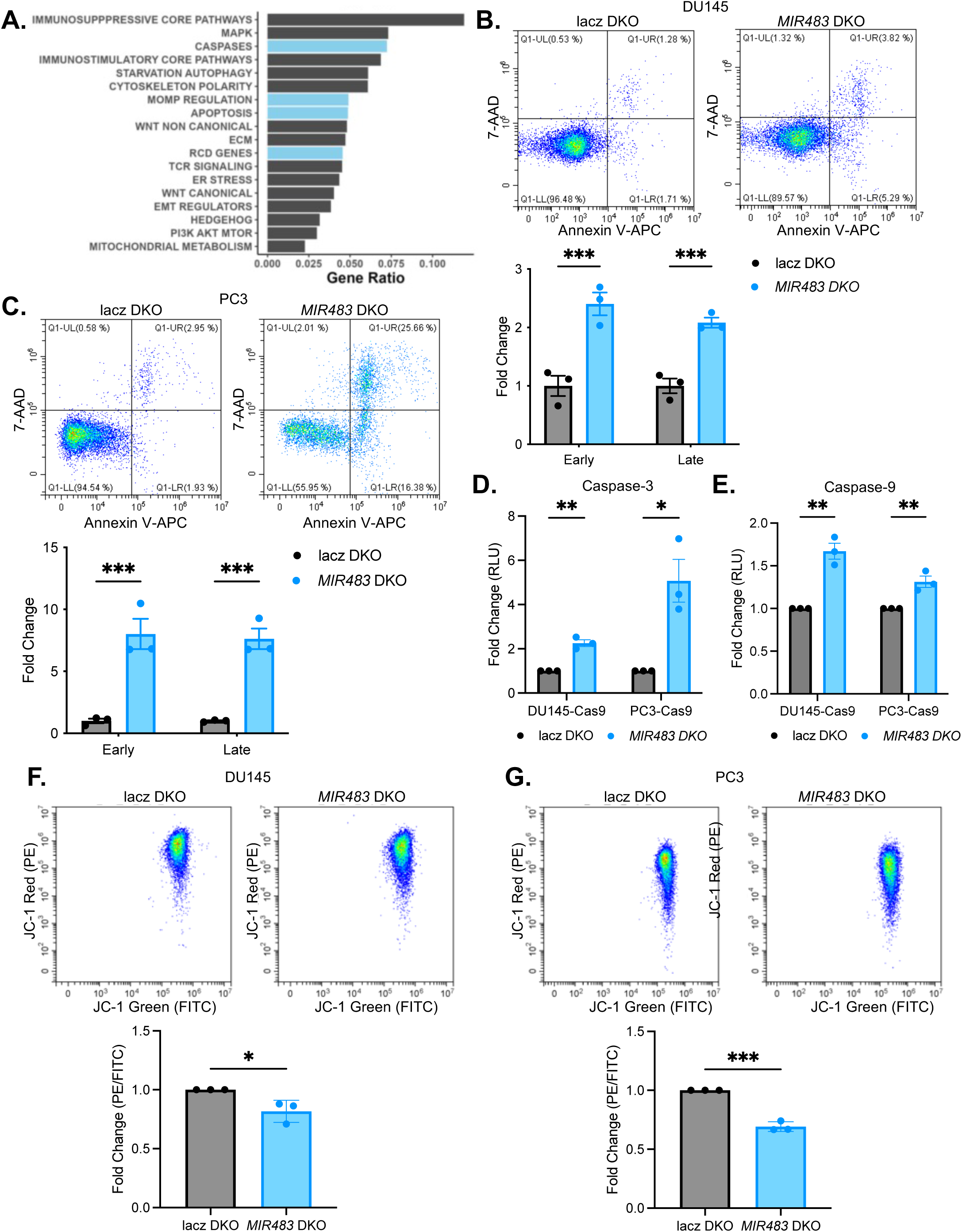
*MIR483* knockout increases apoptosis in PCa cells. **A.** Pathway prediction analysis for miR-483-3p and miR-483-5p. **B.** Annexin V/7-AAD flow cytometry of *MIR483* DKO in DU145-Cas9 cells. Representative plots are shown on top and quantitation is shown on the bottom. **C.** Same as (**B**) in PC3-Cas9 cells. **D.** Caspase-3 activity following *MIR483* DKO in DU145-Cas9 cells (left) and PC3-Cas9 cells (right). **E.** Same as (**D**) with caspase-9 activity. **F.** JC-1 flow cytometry following *MIR483* DKO in DU145-Cas9 cells. Representative plots are shown on the top and quantitation is shown on the bottom. **G.** Same as (**F**) in PC3-Cas9 cells. All data represented as mean ± SEM from n=3 independent experiments. P-values obtained using a two-way ANOVA with a Holm-Sidak’s multiple comparisons test (**B, C**) or an Unpaired one-tailed Student’s t-test (**D-G**). * *p* < 0.05; ** *p* < 0.01; *** *p* < 0.001. See also Figure S3.

Reasoning that decreased cell growth in our KO models was a result of elevated apoptosis, we directly measured various apoptosis parameters experimentally. Firstly, *MIR483* DKO resulted in increased AnnexinV and 7AAD staining in both DU145 and PC3 models (Figure 3B,C).

Secondly, we observed significant upregulation of caspase-3 and caspase-9 activity in both DKO models (Figure 3D,E). Given that caspase-9 is the key executioner caspase in the mitochondrially activated intrinsic apoptotic pathway (34), we evaluated intrinsic apoptosis, specifically mitochondrial outer membrane permeabilization (MOMP), using JC-1, a membrane-permeable cationic dye used to assess mitochondrial membrane potential. *MIR483* DKO in DU145 and PC3 cells was accompanied by a significant decrease in mitochondrial membrane potential (Figures 3F,G and S3A,B). Together these data suggest that loss of miR-483 triggers loss of mitochondrial membrane potential and apoptosis. These results support the notion that miR-483 controls apoptosis through the intrinsic pathway, by regulating mitochondrial homeostasis.

### miR-483-3p targets a BCLAF1/PUMA/BAK1 signaling network to suppress intrinsic apoptosis

We consistently observed that miR-483-3p was expressed at higher levels than miR-483-5p in our PCa cell line panel (Figure S5A) and in patients from the CPC-GENE dataset (Figure S5B) suggesting that miR-483-3p may be the dominant mature strand derived from the miR-483 pre-miRNA hairpin. Building upon previous reports in other malignancies demonstrating that miR-483-3p can suppress apoptosis (35–38), we aimed to elucidate its specific apoptotic role within the context of PCa. By employing the miRNA Data Integration Portal (mirDIP) (39), we identified 72 miR-483-3p targets associated with MOMP (Figure 4A). Given the crucial role of the Bcl-2 family in regulating intrinsic apoptosis and MOMP (40,41), we chose to specifically investigate miR-483-3p targeted Bcl-2 family genes. Of the 7 miR-483-3p-targeted Bcl-2 family genes, four are pro-apoptotic factors, supporting a role for miR-483-3p in suppressing apoptosis (Figure 4A). Among these targets, PUMA and BAK1 emerged as strong candidates targeted by miR-483-3p in PCa cells. PUMA, a known direct target of miR-483-3p in Wilms tumour and neuroblastoma (35,37), demonstrated de-repression in *MIR483* DKO models (Figures 4B,C). Similarly, BAK1 and BAX were de-repressed in *MIR483* DKO models (Figures 4B,C and S5C). Notably, there was no detectable BAX protein expression in DU145 cells (data not shown), similar to previous reports (42). BMF has been reported to have limited pro-apoptotic potential (43), thus it was not studied further.

**Figure 4.**
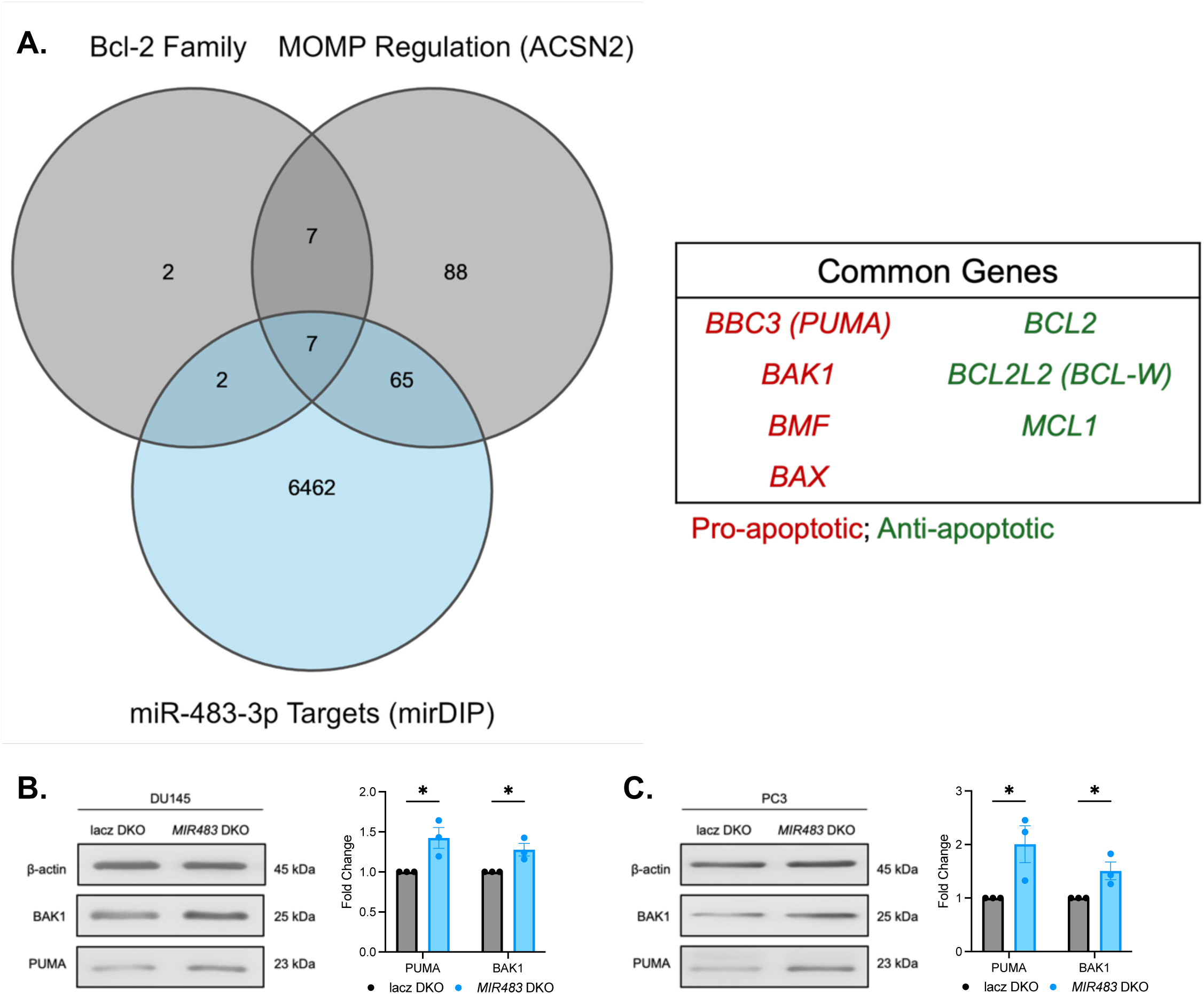
miR-483 is predicted to target apoptosis related genes. **A.** Overlap between predicted targets of miR-483-3p, MOMP-associated genes and Bcl-2 Family genes (left). The 7 common genes are listed on the right. **B.** Western blot of PUMA and BAK1 in *MIR483* DKO DU145-Cas9 cells. Representative blot is shown on the left and quantitation is shown on the right. **C.** Same as (**B**) in PC3-Cas9 cells. All data represented as mean ± SEM from n=3 independent experiments. P-values obtained using an Unpaired one-tailed Student’s t-test. * *p* < 0.05. See also Figure S4 and Figure S5.

Building upon the miRNA predictions, we aimed to confirm that PUMA and BAK1 were direct targets of miR-483-3p. For this, 3’UTR regions of *PUMA* and *BAK1* transcripts containing predicted binding suites were cloned into psiCHECK-2 and tested in a dual luciferase reporter assay. To overexpress miR-483, its genomic locus containing pre-miR-483 plus 100 nucleotides of flanking sequence was cloned downstream of a *U6* promoter in the lentiMIR-puro. Overexpression of both miR-483-3p and miR-483-5p from this construct was confirmed (Figure 5A). We confirmed that PUMA-3’UTR was a direct target of miR-483-3p with a seed-mutant construct which rescued targeting (Figure 5B). Surprisingly, direct targeting of BAK1 was not observed suggesting a more complex mechanism at play (Figure 5B). We hypothesized that BAK1 may be an indirect target of miR-483-3p, therefore we searched for potential regulatory genes of BAK1, such as transcription factors.

**Figure 5.**
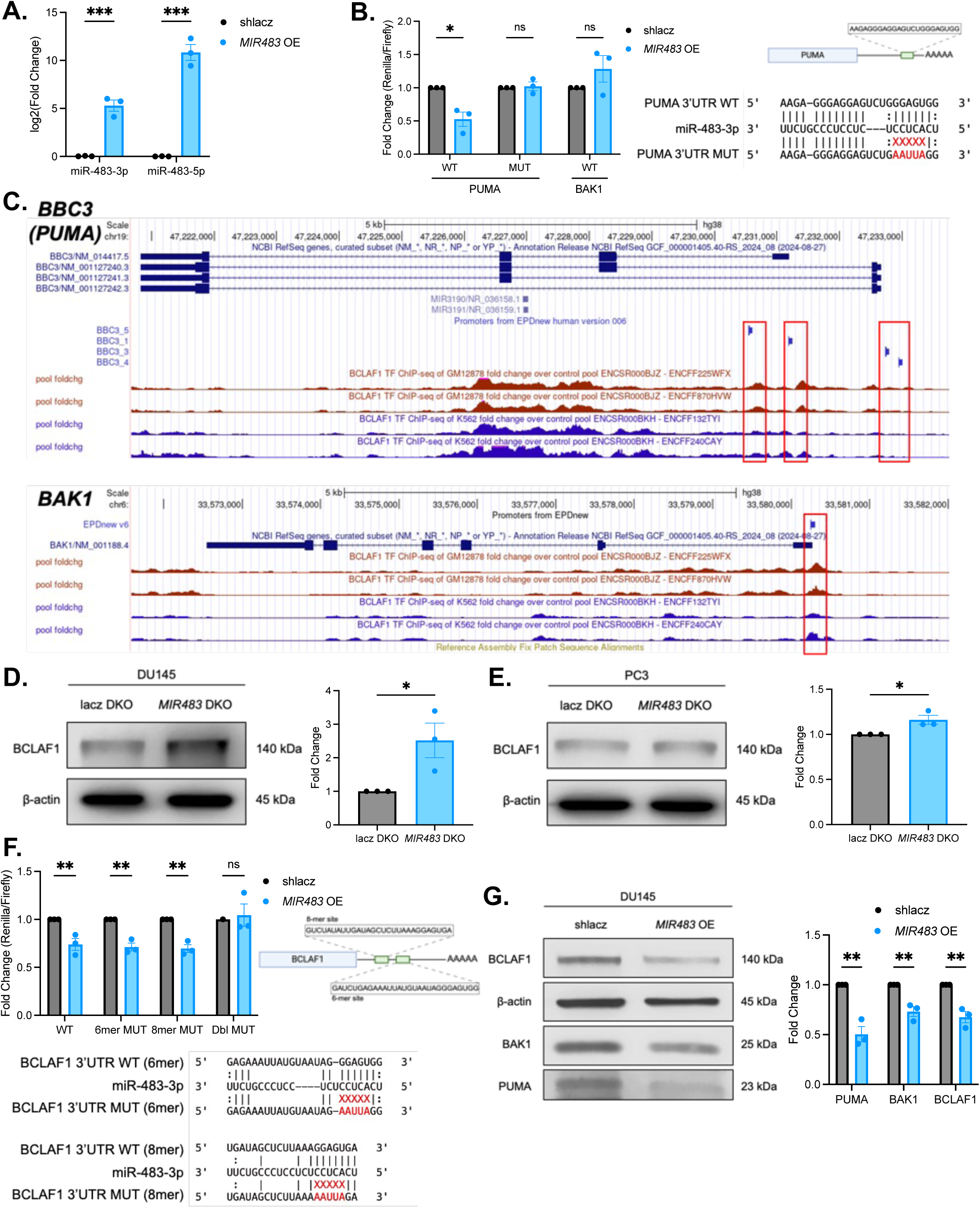
miR-483-3p suppresses a BCLAF1/PUMA/BAK1 signaling network. **A.** RT-qPCR of miR-483-3p and miR-483-5p following *MIR483* overexpression in DU145 cells. **B.** Dual luciferase assay of wild-type (WT) and seed-mutant (MUT) PUMA 3’UTR (left and middle), and WT BAK1 3’UTR reporters in DU145 cells. Schematic representation of the miR-483-3p binding site in the PUMA 3’UTR (top right) and the miR-483-3p seed mutation (bottom right). **C.** Schematic representation of ENCODE ChIPseq peaks and the promoter regions of *BBC3* (top) and *BAK1* (bottom) from GM12878 cells (red track; GSE105550) and K562 cells (blue track; GSE105733). Alignment of ChIPseq peaks and promoter regions is highlighted in red. **D.** Western blot of BCLAF1 in *MIR483* DKO DU145-Cas9 cells. Representative blot is shown on the left and quantitation is shown on the right. **E.** Same as (**D**) in PC3-Cas9 cells. **F.** Same as (**B**) with WT, 6-mer MUT, 8-mer MUT and double MUT BCLAF1 3’UTR reporter. Schematic representation of the miR-483-3p binding sites in the BCLAF1 3’UTR (top right) and the miR-483-3p seed mutations (bottom). **G.** Western blot of BCLAF1, PUMA and BAK1 in *MIR483* overexpressing DU145 cells. Representative blot is shown on the left and quantitation is shown on the right. All data represented as mean ± SEM from n=3 independent experiments. P-values obtained using an Unpaired one-tailed Student’s t-test (**A, D, E, G**) or a two-way ANOVA with a Holm-Sidak’s multiple comparisons test (**B, F**). * *p* < 0.05; ** *p* < 0.01; *** *p* < 0.001; ns, not significant.

By leveraging publicly available transcription factor ChIP-seq data from the ENCODE project (44), we identified that *BAK1* and *BBC3* (encoding PUMA) are putative transcription targets of BCL2 Associated Transcription Factor 1 (BCLAF1) (Figure 5C). Indeed, BCLAF1 has been reported to regulate several apoptotic genes, and promote apoptosis in various cancers (45). In support of this, we observed a significant enrichment of BCLAF1 transcriptional target genes following *MIR483* knockout in MCF7 and PC9 cells (Figure S6). This unexpected finding provides compelling evidence that BCLAF1 was associated miR-483-3p, potentially mediating some of the observed apoptotic effects. Upon examining BCLAF1 protein levels, we observed that it was de-repressed upon *MIR483* DKO in DU145 and PC3 cells (Figures 5D,E). By examining the *BCLAF1* 3’UTR we identified two putative miR-483-3p binding sites, one with a 6-mer seed and a second with an 8-mer seed. We demonstrated that miR-483 overexpression suppressed *BCLAF1* 3’UTR reporter activity, providing further evidence that BCLAF1 is a direct target of miR-483-3p. Peculiarly, single mutation of either the 6-mer seed or 8-mer seed sites was not sufficient to rescue the reporter repression, whereas mutation of both sites did rescue the suppression, confirming that BCLAF1 is a target of miR-483-3p (Figure 5F). Supporting evidence of a role for miR-483-3p in regulating each of BCLAF1, PUMA and BAK1 was also observed upon miR-483 overexpression in DU145 cells (Figure 5G), further indicating the existence of a miR-483-3p/BCLAF1/PUMA/BAK1 signaling network.

### miR-483-3p inhibition increases docetaxel efficacy

We next sought to validate that specific inhibition of miR-483-3p alone, rather than complete disruption of the miR-483 duplex with CRISPR, could confer essentiality phenotypes. For this we utilized Tough Decoys (TD); synthetic RNA molecules engineered to inhibit mature miRNA function by acting as high-affinity sponges or traps (Figure 6A) (46,47). After confirming the efficacy of miR-483-3p TD using a specific miR-483-3p sensor (Figure 6B), we also show that miR-483-3p TD resulted in a significant reduction in PC3 cell growth compared to TD controls (Figure 6C). These data indicated that specific inhibition of miR-483-3p is essential for PCa cell fitness.

**Figure 6.**
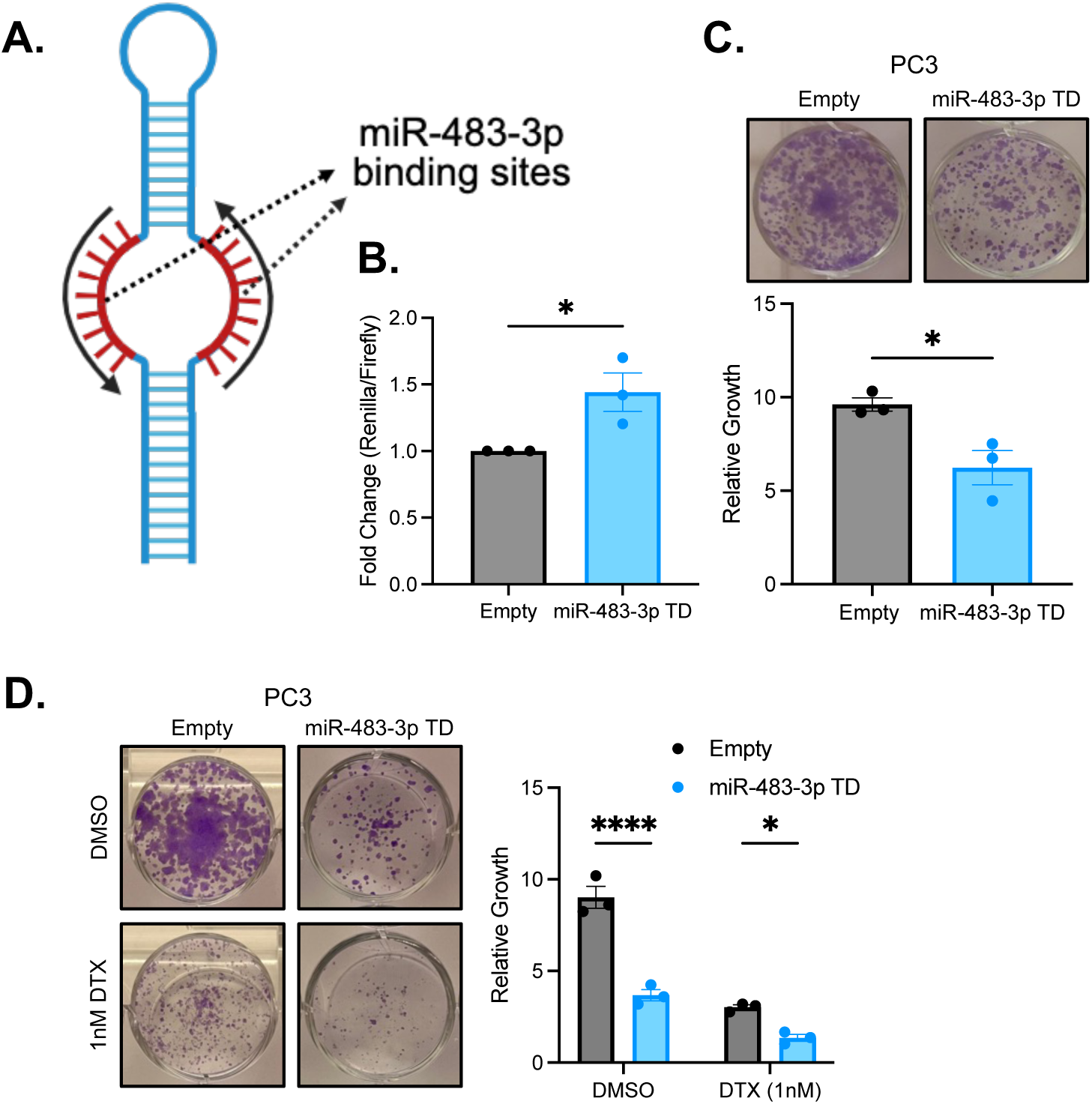
miR-483-3p is essential and sensitizes PCa cells to docetaxel. **A.** Schematic representation of the miR-483-3p TD inhibitor. **B.** Dual luciferase assay of miR-483-3p TD with a specific miR-483-3p sensor in PC3 cells. **C.** Growth assay of miR-483-TD in PC3 cells. Representative images are shown on top and quantitation is shown on the bottom. **D.** Growth assay of miR-483-3p TD in PC3 cells treated with either DMSO or 1nM DTX. Representative images are shown on the left and quantitation are shown on the right. All data represented as mean ± SEM from n=3 independent experiments. P-values obtained using an Unpaired one-tailed Student’s t-test (**B,C**) or a two-way ANOVA with a Holm-Sidak’s multiple comparisons test (**D**). * *p* < 0.05; **** *p* < 0.0001.

Next, the potential of miR-483-3p inhibition to improve chemotherapy efficacy was explored by examining the consequences of combining miR-483-3p TD with docetaxel (DTX), a chemotherapeutic agent used for metastatic PCa (2–4). Our results demonstrate that combined miR-483-3p inhibition with DTX significantly reduced PC3 cell growth compared to either treatment alone (Figure 6D) suggesting that combining miR-483-3p inhibition with apoptosis-inducing chemotherapy, such as DTX, may offer improved treatment outcomes for metastatic PCa.

### miR-483 overexpression promotes growth in premalignant cells but not in PCa cells and attenuates docetaxel efficacy

To investigate the growth promoting potential of miR-483 in PCa cells, we overexpressed miR-483 in DU145 cells. After confirming overexpression of both miR-483-3p and miR-483-5p mature miRNA strands (Figure 5A) we failed to observe any growth difference (Figure 7A). Surprisingly, miR-483 overexpressing cells were resistant to DTX cytotoxicity, suggesting that miR-483 may suppress apoptotic mechanisms normally engaged by DTX (Figure 7B).

**Figure 7.**
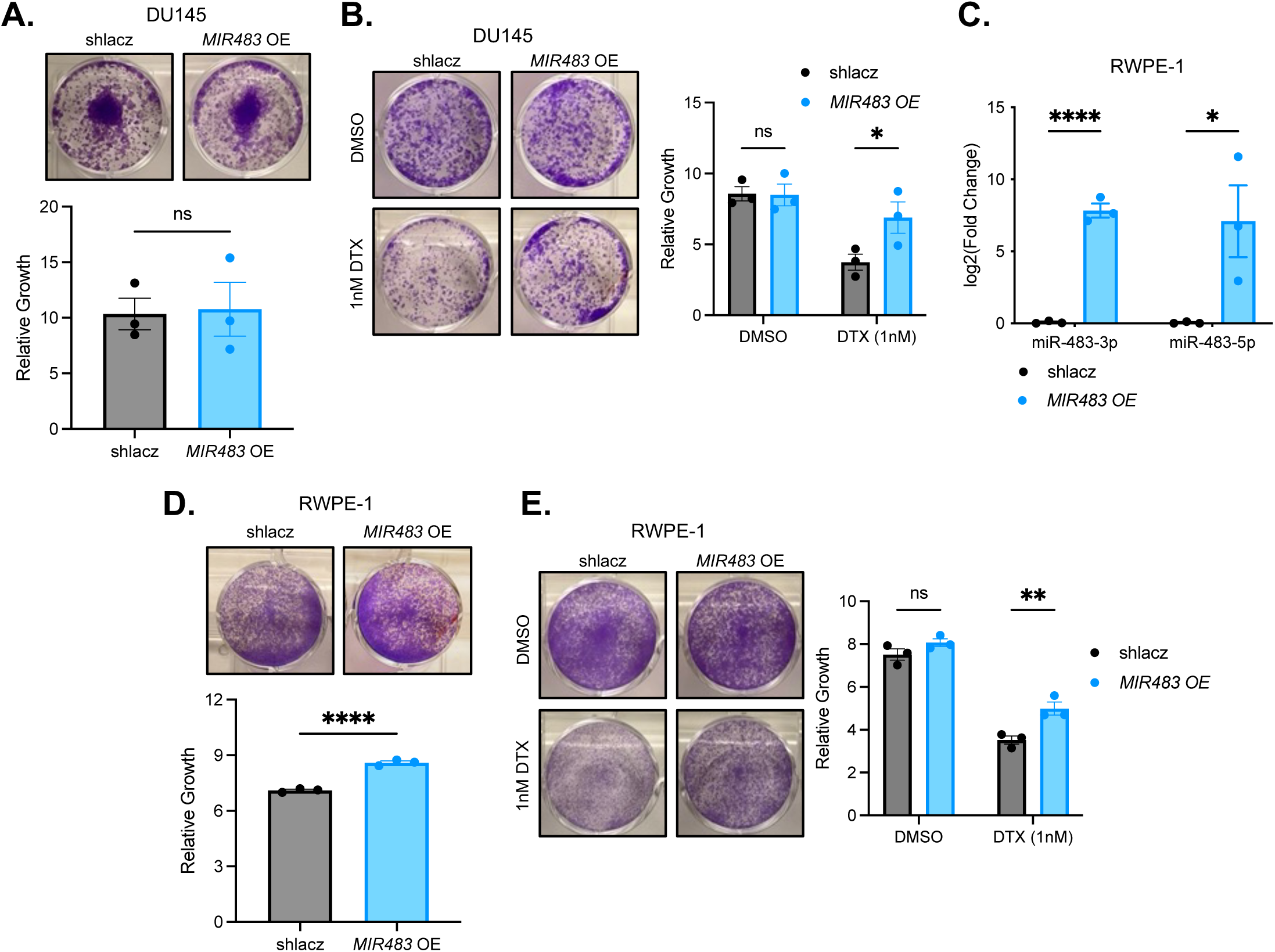
miR-483 promotes growth in premalignant cells but not PCa cells and attenuates docetaxel efficacy. **A.** Growth assay of *MIR483* overexpression in DU145 cells. Representative images are shown on top and quantitation is shown on the bottom. **B.** Growth assay of *MIR483* overexpression in DU145 cells treated with either DMSO or 1nM DTX. Representative images are shown on the left and quantitation are shown on the right. **C.** RT-qPCR of miR-483-3p and miR-483-5p following *MIR483* overexpression in RWPE-1 cells. **D.** Same as (**A**) in RWPE-1 cells. **E.** Same as (**B**) in RWPE-1 cells. All data represented as mean ± SEM from n=3 independent experiments. P-values obtained using an Unpaired one-tailed Student’s t-test (**A,D**) or a two-way ANOVA with a Holm-Sidak’s multiple comparisons test (**B,C,E**). * *p* < 0.05; ** *p* < 0.01; **** *p* < 0.0001; ns, not significant.

**Figure 8.**
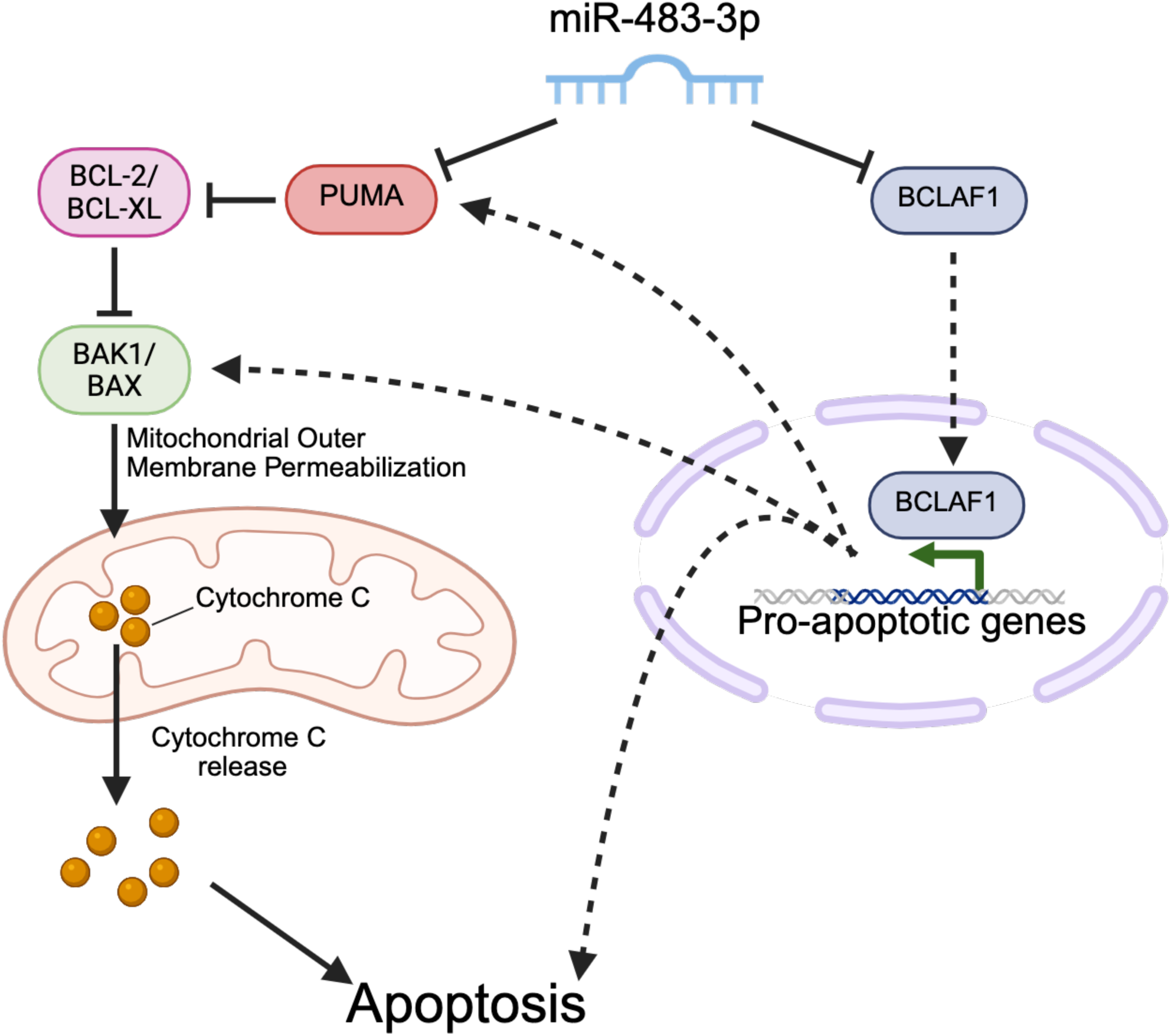
Schematic representation of proposed miR-483-3p mechanism of essentiality. miR-483-3p is essential for PCa cell survival through suppression of apoptosis by targeting a BCLAF1/PUMA/BAK1 network. Inhibiting miR-483-3p in combination with apoptosis inducing agents can have additive effects at reducing PCa cell growth.

To further explore growth promoting functions of miR-483, we overexpressed miR-483 in premalignant RWPE-1 cells (Figure 7C). Unlike DU145 cells, we found that miR-483 overexpression promoted growth in RWPE-1 cells compared to control (Figure 7D). Notably, miR-483 overexpression also conferred DTX resistance in RWPE-1 cells (Figure 7E). Together these results suggest that miR-483 can promote growth in non-transformed cells, however PCa cells are insensitive to elevated miR-483 levels.

## DISCUSSION

In this study we generated miRKOv2: a novel CRISPR library targeting exclusively miRNA to screen PCa cell lines. Compared to previous iterations of miRNA KO libraries, miRKOv2 offers superior on-target efficacy, reduced off-target effects and inclusion of positive control gRNA, permitting the use of the BAGEL algorithm for improved analysis of CRISPR dropout screens (26). We expect miRKOv2 to be a powerful resource to discover high-confidence miRNA candidates responsible for an array of phenotypes. miRKOv2 allowed us to identify essential roles for several miRNA in cellular growth and fitness, challenging the notion that miRNA function solely as fine-tuners of gene expression. Among the essential miRNA, our findings establish miR-483-3p as a master regulator of PCa cell survival, acting through direct modulation of the BCLAF1/PUMA/BAK1 apoptotic signaling network.

Previous studies have implicated miR-483-5p, the duplex partner of miR-483-3p, in promoting PCa growth. miR-483-5p overexpression was demonstrated to promote tumor growth in PCa cells by suppressing RBM5, a well-known tumor suppressor gene that modulates apoptosis by regulating the alternative splicing of several critical apoptotic mediators including FAS and caspase-2 (48–50). In a separate study, miR-483-5p was described as a crucial miRNA in a ceRNA network involving *LINC00908* and the tumour suppressor TSPYL5 which suppresses apoptosis to promote PCa progression (51). When taken together with our findings descried herein, it appears that both the 3p and 5p mature miRNA strands of miR-483 regulate key features of PCa cells involved in disease progression. In our study, unlike miR-483-5p, miR-483-3p was unable to promote PCa cell growth; however, its disruption was crucial for growth. In sum, miR-483-3p fits a gene profile characterized as essential for growth but does not demonstrate oncogenic features when overexpressed in PCa cells.

Nonetheless, its essentiality positions miR-483-3p as a putative therapeutic target in PCa. Notably, miR-483 disruption in premalignant prostate epithelial cells did not result in any growth defect, a finding consistent with a previous study demonstrating that *MIR483* knockout mice developed normally and remained viable (52). This data indicates the existence of a therapeutic window for targeting miR-483 in metastatic PCa, whilst sparing non-transformed tissues.

Based on the observed essentiality of miR-483 in PCa cell survival, we hypothesized that targeting miR-483-3p could render tumor cells more susceptible to apoptosis-inducing drugs. Upon examining the effect of combining miR-483-3p inhibition with docetaxel (DTX) - a standard therapy for metastatic PCa (2–4)-we observed a significant additive effect. These findings suggest that the observed sensitization may be further enhanced through co-treatment with agents such as venetoclax, a Bcl-2 inhibitor currently undergoing clinical evaluation for metastatic PCa. (NCT03751436) (53).

To elucidate the mechanisms underlying miR-483 function in PCa we performed bioinformatic analyses using publicly available datasets to identify candidate pathways associated with miR-483-mediated cell survival. These analyses revealed a significant association between miR-483 and apoptosis-related genes, with Bcl-2 family members—PUMA, BMF, BAK1, and BAX—emerging as key potential mediators. Subsequent experimental validation in PCa cell lines demonstrated that disruption of miR-483 modulates the expression of PUMA and BAK1; however, only PUMA was confirmed as a direct target of miR-483-3p. Crucially, we discovered that BCLAF1, a crucial regulator of both *PUMA* and *BAK1* expression, is also a direct miR-483-3p target (45). This revealed a negative feed-forward loop, wherein miR-483-3p directly and indirectly suppresses Bcl-2 family genes via BCLAF1 (Figure 7), enhancing its apoptotic regulatory capacity (8). Our observation of a miR-483-3p/BCLAF1 feed-forward loop aligns with comparable regulatory networks in PCa, including miR-125b/p53, miR-17/92/MYC/E2F1, and miR-145/IGF1R, all described by Afshar *et al.* (*54*), which underscore the essential nature of miR-483 in PCa cells.

In sum, our study addresses the critical need for robust, large-scale methodologies to investigate miRNA function, particularly in metastatic PCa, a disease with limited treatment options and significant miRNA involvement (55). Specifically, we identified *MIR483* as a novel essential gene in PCa, acting through the BCLAF1/PUMA/BAK1 apoptotic network (Figure 7). The selective vulnerability of PCa cells to miR-483 targeting, coupled with enhanced DTX sensitivity and sparing of normal non-transformed prostate epithelial cells, underscores its potential as a novel therapeutic strategy for metastatic PCa. Collectively this data positions miR-483-3p as a promising therapeutic target for further exploration.

## MATERIALS AND METHODS

### Cell lines and drug treatments

DU145, PC3, LNCaP-FGC (LNCaP) and 22Rv1 cells (ATCC) were cultured in RPMI-1640 medium (Wisent Inc., 350-000-CL) supplemented with 10% fetal bovine serum (FBS; Wisent Inc., 080150) and 1% penicillin/streptomycin (Wisent Inc., 450-201-EL). HEK293T cells (ATCC) were cultured in DMEM medium (Wisent Inc., 319-005-CL) supplemented with 10% FBS and 1% penicillin/streptomycin. RWPE-1 cells (ATCC) were cultured in K-SFM medium supplemented with bovine pituitary extract and recombinant epidermal growth factor (Gibco, 17005042). Cas9^+^ DU145 and Cas9^+^ PC3 cells were generated through viral transduction with lentiCas9-Blast lentivirus and used for all knockout experiments. All cells were maintained at 37°C with 5% CO_2_. DU145, PC3 and RWPE-1 cells were dosed with 1nM docetaxel (CST, 9886S) for the appropriate assays.

### Plasmids and viral vectors

The miRKOv2 library was cloned into a modified version of the lentiGuide-puro vector (Addgene #52963) containing a turboGFP cassette (lentiGuide-puro-P2A-GFP). Constitutive Cas9^+^ cell lines were made using the lentiCas9-Blast vector (Addgene #52962). Single guide RNAs (sgRNAs) targeting *MIR483*, or a double knockout (DKO) system employing two *MIR483* flanking sgRNAs were cloned into the lentiGuide-puro vector or the lentiCRISPRv2 (Addgene #52961) vector and used for validation experiments. For overexpression studies, the *MIR483* genomic locus with 100bp of surrounding context and slack negative control sequences were cloned into the lentiGuide-puro vector. Tough Decoy sequences were cloned into the pLKO.1-puro vector (Addgene #8453) downstream of a *U6* promoter sequence. A sequence of four tandem repeat miR-483-3p binding sites from LSB-hsa-miR-483-3p (Addgene #103550) and all 3’UTR sequences were cloned into the psiCHECK-2 vector (Promega, C8021). Lentivirus was made using the pMDG.2 (Addgene #12259) and psPAX2 (Addgene #12260) vectors.

### Plasmid Cloning

#### Single gRNA cloning

DNA oligos for single gRNA were annealed and cloned into the lentiGuide-puro or lentiCRISPRv2 vectors as described in Shalem *et al.* (56). Single gRNA sequences are listed in Table S2.

#### DKO system cloning

A sequence containing the DKO system (gRNA-tracrRNA-U6-gRNA) was synthesized with flanking BsmBI digestion sites (IDT) (Table S3). This sequence was cloned into the lentiGuide-puro vector similar to the single gRNA cloning described above and Shalem *et al.* (56).

#### Generation of MIR483 overexpression constructs

The *MIR483* genomic locus with 100 nucleotides up- and downstream followed by a *U6* poly(T) termination sequence was synthesized with flanking BsmBI digestion sites (IDT) (Table S3) and cloned into the lentiGuide-puro vector (Addgene). A negative control, lacz-targeting shRNA sequence was also synthesized (IDT) (Table S3) and cloned in a similar manner. We term this overexpression system the lentiMIR-puro system.

#### Tough Decoy inhibitor cloning

The Tough Decoy sequence for miR-483-3p was designed as described in Yoo *et al.* (*46*) and synthesized (IDT) (Table S3). The synthesized sequence was then amplified using primers containing AgeI and EcoRI digestion sites listed in Table S1 and cloned into the pLKO.1 vector.

#### miR-483-3p sensor cloning

A sequence of four tandem repeat miR-483-3p binding sites was amplified from LSB-hsa-miR-483-3p (Addgene) using primers listed in Table S1. The amplified sequence was cloned into the psiCHECK-2 vector (Promega) through Gibson cloning with the NEBuilder HiFi DNA Assembly Master Mix (NEB, E261L).

### Lentiviral production and transduction

9×10^6^ HEK293T cells were seeded in 15cm dishes and co-transfected with 50ug plasmid of interest, 30ug psPAX2 and 15ug pMDG.2 using the calcium phosphate method. Virus was concentrated through ultracentrifugation at 22,000rpm for 2.5 hours, resuspended in 50uL PBS (Wisent Inc., 311-010-CL) and stored at −80°C for later use. Target cells were transduced with lentivirus and supplemented with 8ug/mL protamine sulfate (MP Bioomedicals, 02194729-CF) for 48 hours prior to a changing of medium. 24 hours later, cells were selected with puromycin (2ug/mL for 2 days; BioShop, PUR333) or blasticidin (750ug/mL for 7 days; BioShop, BLA477) depending on the vector.

### Plasmid transfection

3×10^5^ cells (DU145) or 6×10^5^ cells (RWPE-1) were seeded in 6-well plates, or 1×10^5^ cells were seeded in 12-well plates (PC3, DU145) 24 hours prior to transfection. All plasmids were transfected using the Lipofectamine 2000 reagent (Invitrogen, 11668019) according to the manufacturer’s protocol and in antibiotic-free medium. Transfected cells were collected for their appropriate assays 24 hours post-transfection. For miR-483 overexpression, 2ug (6-well) or 1ug (12-well) of each plasmid was transfected. For dual-luciferase sensor constructs, 500ng (12-well) of each plasmid was transfected.

### miRKOv2 design and cloning

#### In silico design

All annotated pre-miRNA and the corresponding sequences were obtained from miRBase v22.2 (57). HGNC gene symbols for each pre-miRNA were used as input for the Broad GPP sgRNA Designer (19) (https://portals.broadinstitute.org/gpp/public/analysis-tools/sgrna-design) to identify all possible gRNA. Next, gRNA were excluded based on on-target score (Rule Set 2 score < 0.2) and off-target score (CFD Match Bin I score > 3) (19). Lastly, gRNA targeting non-loop regions of the pre-miRNA sequence were prioritized for selection. For every pre-miRNA locus, where possible 4 gRNA from the remaining pool were selected with a minimum of 3 gRNA required. Pre-miRNA that did not satisfy this requirement were excluded as targets in the final library. Core-essential and non-essential genes were selected based on Bayes Factor across previous CRISPR-Cas9 screens (21–23), where 100 core-essential genes with the largest Bayes Factor were chosen and 100 non-essential genes with smallest Bayes Factor were chosen. gRNA targeting these core-essential and non-essential genes were identified as described above and selected based on the combined ranking generated by the sgRNA Designer such that each gene was targeted by 4 gRNA. 74 gRNA (1% of the final library) were randomly selected from a pooled list of non-targeting gRNA from previously published libraries (19,58,59). The final miRKOv2 library contains 6532 gRNA targeting 1649 pre-miRNA loci, 400 gRNA targeting 100 core-essential genes, 400 gRNA targeting 100 non-essential genes, and 74 non-targeting gRNA.

#### Benchmarking analysis

To benchmark the miRKOv2 library against existing libraries, we sought to compare the on-target and off-target score profiles. Both the miRKOv2 library and the lentiG-mir library (24) utilize the Rule Set 2/Azimuth 2.0 algorithm for on-target scoring and the Cutting Frequency Determination (CFD) algorithm for off-target scores (19). Since the Lx-miR library (25) and the miRKO library (16) used older on-target and off-target algorithms, we calculated these scores using the Rule Set 2/Azimuth 2.0 and the CFD algorithms. For on-target benchmarking, the distribution of on-target scores was compared between libraries. For off-target benchmarking, the cumulative fraction of loci with a CFD score > 0.2 in a protein-coding gene was compared between libraries.

#### miRKOv2 library amplification and cloning

Selected gRNA for the miRKOv2 library were synthesized as a 73mer pooled oligo array (Custom Array) and amplified using primers listed in Table S1. The amplified oligo pool was cloned into the lentiGuide-puro-P2A-GFP vector through Gibson cloning (QuantiBio, 95190). The resulting Gibson reaction product was transformed into 25uL electrocompetent cells (Endura, 60242) according to the manufacturer’s protocol and plated onto 8 LB-agar plates in 245mm^2^ BioAssay dishes (Corning, 431111). Plasmid DNA from the resulting colonies were extracted with the QIAGEN Plasmid Maxi Kit (Qiagen, 12163).

### miRKOv2 Dropout Screen

2.5×10^7^ DU145 or LNCaP cells were transduced with miRKOv2 lentivirus at an MOI of 0.3 and at a ∼1000X gRNA coverage (∼7.5×10^6^ cells) and selected with puromycin as described above. After puromycin selection (T0), the remaining pool of cells was split into 3 technical replicates, each maintained at a 1000X gRNA coverage. Remaining cells were harvested for genomic DNA extraction. Each screen replicate was passaged every 3 days for a total of 27 days (T3-T27) or every 6 days for a total of 30 days (T6–T30) for DU145 and LNCaP, respectively. Remaining cells at each time point were collected for genomic DNA extraction.

### Screen sequencing and analysis

Genomic DNA was extracted from collected cells at each time point using the QIAamp DNA Blood Midi Kit (Qiagen, 51185) according to the manufacturer’s protocol. Samples were submitted to the Princess Margaret Genomics Centre (PMGC, University Health Network, Toronto, Canada) for NGS library preparation and sequencing. Samples were sequenced at a depth corresponding to ∼500X gRNA coverage. Read counts from all timepoints can be found in File S3 for the DU145-Cas9 dropout screen or in File S4 for the LNCaP dropout screen.

Demultiplexed and trimmed reads were aligned to the miRKOv2 library using Bowtie2 tool (60). After alignment, read counts were calculated and essential miRNA were identified using the Bayesian Analysis of Gene Essentiality (BAGEL) algorithm, as described in Hart and Moffat (2016) (26) and the reference set of gRNA targeting core-essential genes (n=400) and non-essential genes (n=400) (21–23) that were included in the miRKOv2 library design and using a supervised iterative learning method (n=1000 iterations). A gene was considered to be essential if it had a BF > 0 and an FDR < 0.1.

### Growth Assays

For 6 day growth assays, 1×10^3^ cells (PC3) or 5×10^3^ cells (DU145) were seeded in 12-well plates. For 10 day growth assays, 1×10^3^ cells (DU145) were seeded in 12-well plates. 24 hours later and at the terminal timpoint, cells were fixed with 10% formalin (Sigma Aldrich, HT501128-4L) prior to staining with 0.05% crystal violet dissolved in 40% v/v methanol and destaining with water.

Stained plates were imaged and dissolved with 10% acetic acid for quantitation of absorption at 595nm using the SpectraMax M3 using the SoftMaxPro v6 software (Molecular Devices).

### Protein extraction and Western Blot

Whole cell protein lysates were extracted from cells using 1X RIPA buffer (CST, 9806) supplemented with protease inhibitors (Roche, 11836163001) and sonicated at 30% amplitude. Blots were imaged using the KwikQuant Ultra Digital-ECL Subtrate (Kindle Biosciences, R1002) and the KwikQuant Imager. The following primary antibodies were used in this study: BCLAF1 (1:1000; Invitrogen, PA5-55686), ꞵ-actin (1:10,000; CST, 4967), PUMA (1:1000; CST 12450T), BAK1 (1:1000; CST, 12105S), BAX (1:1000; CST, 5023T), IGF2 (1:1000; Invitrogen, MA5-17096). All original blots can be found in Figure S7.

### RNA extraction and RT-qPCR

Total cellular RNA was extracted using the RNeasy Mini kit (Qiagen, 74104). miRNA cDNA synthesis was performed using the TaqMan Advanced miRNA cDNA Synthesis kit (Applied Biosystems, A28007). Quantitative PCR was performed on cDNA using the TaqMan Fast Advanced Master Mix (Applied Biosystems, 4444557) with TaqMan Advanced miRNA assays (Applied Biosystems, A25576) and the QuantStudio 3 Real-Time PCR system (Applied Biosciences) using the QuantStudio Design and Analysis software v1.2 (Applied Biosciences). All Ct values were normalized to hsa-miR-103a-3p and using the ΔΔCt method. The following TaqMan assays were used in this study: hsa-miR-483-3p (Applied Biosystems, 478122_mir), hsa-miR-483-5p (Applied Biosystems, 478432_mir), hsa-miR-103a-3p (Applied Biosystems, 478453_mir).

### Characterization of indels

The *MIR483* locus with 300 nucleotides up- and down-stream were amplified using primers listed in Table S1 from extracted genomic DNA and submitted to The Centre for Applied Genomics (TCAG, The Hospital for Sick Children, Toronto, Canada) for sanger sequencing with a sequencing primer listed in Table S1. The control and mir-483 DKO sequencing chromatograms were then inputted in the Tracking of Indels by Deconvolution (TIDE) tool (29) (https://tide.nki.nl/) for indel deconvolution and identification.

### miRNA-target interaction analysis

#### Amplification of 3’UTR sequences and cloning into reporter constructs

3’UTR fragments (up to 1.5kb) were amplified from genomic DNA extracted from PC3 cells using primers listed in Table S1 and cloned into the digested psiCHECK-2 vector (Promega) using Gibson cloning with the NEBuilder HiFi DNA Assembly Master Mix (NEB, E261L).

#### Generation of seed mutations

Predicted miRNA binding sites were obtained using the DIANA-micro-T (2023) tool (61). The resulting binding sites were aligned with the cloned 3’UTR reporters. Mutations in the seed region from positions 3-7 (GGAGU > AAUUA) were generated by site directed mutagenesis using mutagenic primers listed in Table S1 and the Q5 Site Directed Mutagenesis kit (NEB, E0554S).

#### Dual luciferase assay and analysis

Target cells were transfected with reporter constructs. 24 hours post-transfection, cells were harvested to perform the dual luciferase assay with the Dual-Glo Luciferase Assay System (Promega, E2920) according to the manufacturer’s protocol in white bottom 96 well plates (Corning, 3917). Luminescence was measured using the SpectraMax M3 using the SoftMaxPro v6 software (Molecular Devices). The Renilla/Firefly ratio for each well was calculated and the ratio fold change was calculated relative to the corresponding negative control samples.

### Flow cytometry and Fluorescence Imaging

#### Annexin V/7-AAD staining

1×10^5^ cells were resuspended in 1X Annexin V binding buffer (Invitrogen, 88-8007-74) prior to incubation with APC-conjugated Annexin V antibody (Invitrogen, 88-8007-74) and 7-AAD (CST, 47501S) for 10 minutes on ice in the dark. Cells were analyzed with the CytoFlex flow cytometer (Beckman Coulter) using the CytExpert software v4 (Beckman Coulter). At least 1×10^4^ single cells were collected per sample. Fluorescence measurements for Annexin V and 7-AAD were detected using the APC and PC5.5 channels, respectively. Early apoptosis was defined as Annexin V^+^/7-AAD^-^ cells and late apoptosis was defined as Annexin V^+^/7-AAD^+^ cells.

#### JC-1 staining

JC-1 detection and analysis was performed as previously described (62). Briefly, 1×10^5^ cells were stained with 1uM JC-1 (CST, 92891S) for 30 minutes at 37°C in the dark prior to analysis by flow cytometry or fluorescence imaging. For flow cytometry, cells were washed with PBS (Wisent) prior to analysis with the CytoFlex flow cytometer (Beckman Coulter) using a 488nm laser and the CytExpert software v4 (Beckman Coulter). At least 1×10^4^ single cells were collected per sample. Fluorescence measurements for JC-1 aggregates and JC-1 monomers were detected using the PC5.5 and FITC channels, respectively. For fluorescence imaging, cells were washed and imaged in PBS at 40X objective using the EVOS FL inverted fluorescence microscope. JC-1 aggregates and JC-1 monomers were imaged using the Texas-Red and GFP filters. Brightness for all images was adjusted to improve fluorescence signal.

### Caspase activity assays

1×10^4^ cells were collected to assess caspase activity using the caspase-3 or caspase-9 glo reagents (Promega, G8090 and G8210 respectively) according to the manufacturer’s protocol in white bottom 96 well plates (Corning). The resulting luminescence was measured with the SpectraMax M3 using the SoftMaxPro v6 software (Molecular Devices).

### miRNA Expression Sequencing and Analysis

Total RNA from DU145, PC3, LNCaP, 22Rv1 and VCaP cells was extracted using the RNeasy Mini kit (Qiagen) prior to submission to the TCAG sequencing facility (The Hospital for Sick Children, Toronto, Canada) for small RNA library and sequencing on the Illumina NovaSeq 6000. The resultant sequence reads were demultiplexed and trimmed using Cutadapt (63). Reads were then aligned and annotated to miRBase v22.2 (57) using the BWA alignment tool (v0.7.17) (64). miRNA were considered to be constitutively expressed if they had ≥ 5 reads in at least 80% of the cell line panel.

### miRNA Expression Analysis in Patient Samples

miRNA expression profiles from patients in the CPC-GENE patient dataset were obtained from GSE135535. The expression profiles of miR-483-3p and miR-483-5p were then extracted and plotted.

### Pathway Analyses

#### Prediction of downstream miRNA targets and pathways

Predicted targets and pathways downstream of mir-483 mature strands were identified using the miDIP target prediction tool (39) and mirDIP-integrated pathDIP v.5 tool (65), respectively. For pathway predictions, the top 1% of predicted targets were selected for pathway enrichment analysis using the pathDIP algorithm (65). The analysis was conducted using literature curated pathway associations related to the Atlas of Cancer Signaling Network v2 (ACSN2) (33). Significantly enriched pathways were identified based on a Bonferroni q < 0.05. The gene ratio was calculated based on the ratio between predicted targets and pathway gene size.

#### Gene set enrichment analysis (GSEA)

GSEA was performed on the GSE242259 (Merk *et al*. (24)) dataset using GSEA v.4.3.0 provided by the Broad Institute (66,67) (https://www.gsea-msigdb.org/gsea/index.jsp). Samples were split by *MIR483* KO or control. Enriched gene sets were identified by 1000 gene set permutations. Gene sets with a nominal *p*-value < 0.05 were considered significantly enriched.

The curated KEGG (CP:KEGG) gene set was obtained from MSigDB Collections (32) (https://www.gsea-msigdb.org/gsea/msigdb/collections.jsp).

The curated ACSN2 gene set was obtained from the Atlas of Cancer Signaling Network (33) (https://acsn.curie.fr/ACSN2/downloads.html).

The BCLAF1 targets gene set was obtained from Harmonizome 3.0 database (68) (https://maayanlab.cloud/Harmonizome/gene_set/BCLAF1/ENCODE+Transcription+Factor+Targets).

### Analysis of Transcription Factor Chromatin Immunoprecipitation Sequencing (TF-ChIPseq) data

BCLAF1 TF-ChIPseq peaks in K562 cells (GSE105733) and GM12878 cells (GSE105550) were obtained from the ENCODE project (44). Promoter regions were obtained and imported from the Eukaryotic Promoter Database (EPD) (69). BCLAF1 TF-ChIPseq peaks and promoter regions for *BBC3 (PUMA)* and *BAK1* were aligned to the GRCh38 reference genome and visualized with the UCSC Genome Browser (70) (https://genome.ucsc.edu).

### Statistical Analysis

All statistical analyses were conducted using GraphPad Prism v9. All data is from 3 independent experiments unless otherwise indicated. Where specified in the experimental section, unpaired one-tailed Student’s t-test was used for comparisons between two groups, and one-way ANOVA or two-way ANOVA tests were used for comparisons between more than two groups where appropriate. Multiple comparisons were calculated using the Holm-Sidak method. *, **, *** and **** denote p < 0.05, < 0.01, < 0.001 and < 0.0001, respectively. n.s. denotes no significance.

## DATA AVAILABILITY

Any additional information and materials in this paper are available from the corresponding author upon reasonable request.

## ACKNOWLEDGMENTS

We would like to thank all past and present members of the Salmena Lab for their contributions. L.S. is recipient of Tier II Canada Research Chair (CRC) and was supported by a Human Frontier Career Development Program (HFSP) award. Funding for this research was provided in part by Temerty Faculty of Medicine and Department of Pharmacology and Toxicology, University of Toronto and awards received from Canada Foundation for Innovation (CFI-33505) and in part from the Canadian Institute of Health Research (CIHR511837). J.T.S.C. is funded by a CIHR Doctoral Research Award (CIHR181364).

## AUTHOR CONTRIBUTIONS

Conceptualization, J.T.S.C., M.M.M. and L.S.; Methodology, J.T.S.C., M.M.M. and L.S.; Investigation, J.T.S.C., A.D., D.K.C.L., I.A.G., S.C., N.J.F., M.M.M. and L.S.; Formal Analysis, J.T.S.C. and L.S.; Writing – Original Draft, J.T.S.C., A.D. and L.S.; Writing – Revision & Editing, J.T.S.C. and L.S.; Funding Acquisition, L.S.

## COMPETING INTERESTS

The authors declare no competing interests.

## SUPPLEMENTAL INFORMATION

File S1. Excel file containing all Bayes Factor scores at the terminal timepoints for the DU145-Cas9 and LNCaP dropout screens. Related to Figure 1.

File S2. Excel file containing expression of the 32 miRNA hits from the DU145-Cas9 dropout screen and 47 miRNA hits from the LNCaP dropout screen in a PCa cell line panel. Related to Figure 2.

File S3. Read count table for every time point in the DU145-Cas9 dropout screen.

File S4. Read count table for every time point in the LNCaP dropout screen.

**Table S1.** Excel file containing primer sequences used in this study.

**Table S2.** Excel file containing the single gRNA sequences used in this study.

**Table S3.** Excel file containing synthesized sequences used in this study.

### Supplementary Figure Legends

**Figure S1. Quality control of the miRKOv2 dropout screen in DU145-Cas9 cells. Related to Figure 1. A.** Precision-Recall analysis of all time points in the DU145-Cas9 dropout screen. Area under the curve (AUC) calculations are shown in the inset. **B.** Same as (**A**) but for the LNCaP dropout screen.

**Figure S2. Validation of *MIR483* as an essential gene in PCa cells. Related to Figure 2. A.** Log_2_ transformed fold-changes of each gRNA from the miRKOv2 library across all timepoints in the DU145-Cas9 screen. **B.** Same as (**A**) but in the LNCaP dropout screen. **C.** Schematic representation of the DKO system where two gRNA target opposite ends of the *MIR483* gene. **D.** Indel characterization of the *MIR483* locus in *MIR483* DKO DU145-Cas9 cells. **E.** Same as (**D**) in PC3-Cas9. **F.** Western blot of IGF2 in *MIR483* DKO DU145-Cas9 cells. Representative blot is shown on the left and quantitation is shown on the right. **G.** Same as (**F**) in PC3-Cas9 cells. All data in (**F,G**) is represented as mean ± SEM from n=3 independent experiments. P-values obtained using an Unpaired one-tailed Student’s t-test. n.s. not significant.

**Figure S3. *MIR483* knockout induces mitochondrial permeabilization. Related to Figure 3. A.** Representative fluorescent images taken of JC-1 stained control and *MIR483* DKO DU145-Cas9 cells. Scale bar represents 100um. **B.** Same as (**A**) in PC3-Cas9 cells. Brightness for all images was adjusted to improve fluorescence signal.

**Figure S4. *MIR483* knockout is associated with apoptotic pathways and negatively associated with mitochondrial pathways.** Related to Figure 4. A. Gene set enrichment analysis of RNAseq of control and *MIR483* knockout MCF7. **B.** Same as (**A**) in PC9 cells. **C.** Same as (**A**) in HT29 cells. All data obtained from the GSE242259 dataset(24).

**Figure S5. Expression of miR-483-3p in DU145 and PC3 cells. Related to Figure 4. A.** Log_2_ transformed normalized expression of miR-483-3p and miR-483-5p in a PCa cell line panel of LNCaP, VCaP, 22Rv1, DU145 and PC3 cells. **B.** Same as (**A**) in the CPC-GENE dataset (GSE135535). **C.** Western blot of BAX in *MIR483* DKO PC3-Cas9 cells. Representative blot is shown on the left and quantitation is shown on the right. All data is represented as mean ± SEM from n=3 independent experiments (**A, C**) or from n=320 patients (**B**). P-values obtained using an Unpaired one-tailed Student’s t-test. ** *p* < 0.01.

**Figure S6. BCLAF1 transcriptional targets are enriched in *MIR483* knockout cells. Related to Figure 5**. Gene set enrichment analysis using a curated gene set of BCLAF1 targets of RNAseq of control and *MIR483* knockout MCF7 and PC9 cells. All data obtained from the GSE242259 dataset.

## Notes

### Competing Interest Statement

The authors have declared no competing interest.

## REFERENCES

1. Bray F, Laversanne M, Sung H, Ferlay J, Siegel RL, Soerjomataram I, et al. Global cancer statistics 2022: GLOBOCAN estimates of incidence and mortality worldwide for 36 cancers in 185 countries. CA Cancer J Clin. 2024 Apr 4;74(3):229–63.

2. Rebello RJ, Oing C, Knudsen KE, Loeb S, Johnson DC, Reiter RE, et al. Prostate cancer. Nat Rev Dis Primers. 2021 Feb 4;7(1):9.

3. Hamid AA, Sayegh N, Tombal B, Hussain M, Sweeney CJ, Graff JN, et al. Metastatic Hormone-Sensitive Prostate Cancer: Toward an Era of Adaptive and Personalized Treatment. Am Soc Clin Oncol Educ Book. 2023 May;43:e390166.

4. Parker C, Castro E, Fizazi K, Heidenreich A, Ost P, Procopio G, et al. Prostate cancer: ESMO Clinical Practice Guidelines for diagnosis, treatment and follow-up. Ann Oncol. 2020 Sep;31(9):1119–34.

5. Gebert LFR, MacRae IJ. Regulation of microRNA function in animals. Nat Rev Mol Cell Biol. 2019 Jan;20(1):21–37.

6. Friedman RC, Farh KK-H, Burge CB, Bartel DP. Most mammalian mRNAs are conserved targets of microRNAs. Genome Res. 2009 Jan;19(1):92–105.

7. Esteller M. Non-coding RNAs in human disease. Nat Rev Genet. 2011 Nov 18;12(12):861–74.

8. Bracken CP, Scott HS, Goodall GJ. A network-biology perspective of microRNA function and dysfunction in cancer. Nat Rev Genet. 2016 Dec;17(12):719–32.

9. Seyhan AA. Trials and tribulations of microrna therapeutics. Int J Mol Sci. 2024 Jan 25;25(3).

10. Ambs S, Prueitt RL, Yi M, Hudson RS, Howe TM, Petrocca F, et al. Genomic profiling of microRNA and messenger RNA reveals deregulated microRNA expression in prostate cancer. Cancer Res. 2008 Aug 1;68(15):6162–70.

11. Chiosea S, Jelezcova E, Chandran U, Acquafondata M, McHale T, Sobol RW, et al. Up-regulation of dicer, a component of the MicroRNA machinery, in prostate adenocarcinoma. Am J Pathol. 2006 Nov;169(5):1812–20.

12. Poliseno L, Salmena L, Riccardi L, Fornari A, Song MS, Hobbs RM, et al. Identification of the miR-106b∼25 microRNA cluster as a proto-oncogenic PTEN-targeting intron that cooperates with its host gene MCM7 in transformation. Sci Signal. 2010 Apr 13;3(117):ra29.

13. Das R, Gregory PA, Fernandes RC, Denis I, Wang Q, Townley SL, et al. MicroRNA-194 Promotes Prostate Cancer Metastasis by Inhibiting SOCS2. Cancer Res. 2017 Feb 15;77(4):1021–34.

14. Mercatelli N, Coppola V, Bonci D, Miele F, Costantini A, Guadagnoli M, et al. The inhibition of the highly expressed miR-221 and miR-222 impairs the growth of prostate carcinoma xenografts in mice. PLoS ONE. 2008 Dec 24;3(12):e4029.

15. Kim K, Kim HH, Lee C-H, Kim S, Cheon GJ, Kang KW, et al. Therapeutic efficacy of modified anti-miR21 in metastatic prostate cancer. Biochem Biophys Res Commun. 2020 Aug 27;529(3):707–13.

16. Gabra M, Pastrello C, Machado N, Chow JT-S, Kotlyar M, Tokar T, et al. Essential gene networks in acute myeloid leukemia identified using a microRNA -knockout CRISPR library screen. BioRxiv. 2019 Jun 6;

17. Campbell KJ, Leung HY. Evasion of cell death: A contributory factor in prostate cancer development and treatment resistance. Cancer Lett. 2021 Nov 1;520:213–21.

18. McKenzie S, Kyprianou N. Apoptosis evasion: the role of survival pathways in prostate cancer progression and therapeutic resistance. J Cell Biochem. 2006 Jan 1;97(1):18–32.

19. Doench JG, Fusi N, Sullender M, Hegde M, Vaimberg EW, Donovan KF, et al. Optimized sgRNA design to maximize activity and minimize off-target effects of CRISPR-Cas9. Nat Biotechnol. 2016 Feb;34(2):184–91.

20. Chang H, Yi B, Ma R, Zhang X, Zhao H, Xi Y. CRISPR/cas9, a novel genomic tool to knock down microRNA in vitro and in vivo. Sci Rep. 2016 Feb 29;6:22312.

21. Hart T, Tong AHY, Chan K, Van Leeuwen J, Seetharaman A, Aregger M, et al. Evaluation and Design of Genome-Wide CRISPR/SpCas9 Knockout Screens. G3 (Bethesda). 2017 Aug 7;7(8):2719–27.

22. Hart T, Chandrashekhar M, Aregger M, Steinhart Z, Brown KR, MacLeod G, et al. High-Resolution CRISPR Screens Reveal Fitness Genes and Genotype-Specific Cancer Liabilities. Cell. 2015 Dec 3;163(6):1515–26.

23. Hart T, Brown KR, Sircoulomb F, Rottapel R, Moffat J. Measuring error rates in genomic perturbation screens: gold standards for human functional genomics. Mol Syst Biol. 2014 Jul 1;10(7):733.

24. Merk DJ, Paul L, Tsiami F, Hohenthanner H, Kouchesfahani GM, Haeusser LA, et al. CRISPR-Cas9 screens reveal common essential miRNAs in human cancer cell lines. Genome Med. 2024 Jun 17;16(1):82.

25. Kurata JS, Lin R-J. MicroRNA-focused CRISPR-Cas9 library screen reveals fitness-associated miRNAs. RNA. 2018 Jul;24(7):966–81.

26. Hart T, Moffat J. BAGEL: a computational framework for identifying essential genes from pooled library screens. BMC Bioinformatics. 2016 Apr 16;17:164.

27. Bello D, Webber MM, Kleinman HK, Wartinger DD, Rhim JS. Androgen responsive adult human prostatic epithelial cell lines immortalized by human papillomavirus 18. Carcinogenesis. 1997 Jun;18(6):1215–23.

28. Kukkonen K, Autio-Kimura B, Rauhala H, Kesseli J, Nykter M, Latonen L, et al. Nonmalignant AR-positive prostate epithelial cells and cancer cells respond differently to androgen. Endocr Relat Cancer. 2022 Dec 1;29(12):717–33.

29. Brinkman EK, Chen T, Amendola M, van Steensel B. Easy quantitative assessment of genome editing by sequence trace decomposition. Nucleic Acids Res. 2014 Dec 16;42(22):e168.

30. Rodriguez A, Griffiths-Jones S, Ashurst JL, Bradley A. Identification of mammalian microRNA host genes and transcription units. Genome Res. 2004 Oct;14(10A):1902–10.

31. Sélénou C, Brioude F, Giabicani E, Sobrier M-L, Netchine I. IGF2: development, genetic and epigenetic abnormalities. Cells. 2022 Jun 10;11(12).

32. Kanehisa M, Goto S. KEGG: Kyoto encyclopedia of genes and genomes. Nucleic Acids Res. 2000 Jan 1;28(1):27–30.

33. Kuperstein I, Bonnet E, Nguyen HA, Cohen D, Viara E, Grieco L, et al. Atlas of Cancer Signalling Network: a systems biology resource for integrative analysis of cancer data with Google Maps. Oncogenesis. 2015 Jul 20;4(7):e160.

34. Budihardjo I, Oliver H, Lutter M, Luo X, Wang X. Biochemical pathways of caspase activation during apoptosis. Annu Rev Cell Dev Biol. 1999;15:269–90.

35. Veronese A, Lupini L, Consiglio J, Visone R, Ferracin M, Fornari F, et al. Oncogenic role of miR-483-3p at the IGF2/483 locus. Cancer Res. 2010 Apr 15;70(8):3140–9.

36. Lupini L, Pepe F, Ferracin M, Braconi C, Callegari E, Pagotto S, et al. Over-expression of the miR-483-3p overcomes the miR-145/TP53 pro-apoptotic loop in hepatocellular carcinoma. Oncotarget. 2016 May 24;7(21):31361–71.

37. Wu K, Wang J, He J, Chen Q, Yang L. miR-483-3p promotes proliferation and migration of neuroblastoma cells by targeting PUMA. Int J Clin Exp Pathol. 2018 Feb 1;11(2):490– 501.

38. Che G, Gao H, Tian J, Hu Q, Xie H, Zhang Y. MicroRNA-483-3p Promotes Proliferation, Migration, and Invasion and Induces Chemoresistance of Wilms’ Tumor Cells. Pediatr Dev Pathol. 2020;23(2):144–51.

39. Tokar T, Pastrello C, Rossos AEM, Abovsky M, Hauschild A-C, Tsay M, et al. mirDIP 4.1-integrative database of human microRNA target predictions. Nucleic Acids Res. 2018 Jan 4;46(D1):D360–70.

40. Kalkavan H, Green DR. MOMP, cell suicide as a BCL-2 family business. Cell Death Differ. 2018 Jan;25(1):46–55.

41. King LE, Hohorst L, García-Sáez AJ. Expanding roles of BCL-2 proteins in apoptosis execution and beyond. J Cell Sci. 2023 Nov 15;136(22).

42. Tamaki H, Harashima N, Hiraki M, Arichi N, Nishimura N, Shiina H, et al. Bcl-2 family inhibition sensitizes human prostate cancer cells to docetaxel and promotes unexpected apoptosis under caspase-9 inhibition. Oncotarget. 2014 Nov 30;5(22):11399–412.

43. Piñon JD, Labi V, Egle A, Villunger A. Bim and Bmf in tissue homeostasis and malignant disease. Oncogene. 2008 Dec;27 Suppl 1:S41–52.

44. ENCODE Project Consortium. An integrated encyclopedia of DNA elements in the human genome. Nature. 2012 Sep 6;489(7414):57–74.

45. Yu Z, Zhu J, Wang H, Li H, Jin X. Function of BCLAF1 in human disease. Oncol Lett. 2022 Feb;23(2):58.

46. Yoo J, Hajjar RJ, Jeong D. Generation of Efficient miRNA Inhibitors Using Tough Decoy Constructs. Methods Mol Biol. 2017;1521:41–53.

47. Haraguchi T, Ozaki Y, Iba H. Vectors expressing efficient RNA decoys achieve the long-term suppression of specific microRNA activity in mammalian cells. Nucleic Acids Res. 2009 Apr;37(6):e43.

48. Yang Z-G, Ma X-D, He Z-H, Guo Y-X. miR-483-5p promotes prostate cancer cell proliferation and invasion by targeting RBM5. Int Braz J Urol. 2017;43(6):1060–7.

49. Bonnal S, Martínez C, Förch P, Bachi A, Wilm M, Valcárcel J. RBM5/Luca-15/H37 regulates Fas alternative splice site pairing after exon definition. Mol Cell. 2008 Oct 10;32(1):81–95.

50. Fushimi K, Ray P, Kar A, Wang L, Sutherland LC, Wu JY. Up-regulation of the proapoptotic caspase 2 splicing isoform by a candidate tumor suppressor, RBM5. Proc Natl Acad Sci USA. 2008 Oct 14;105(41):15708–13.

51. Fan L, Li H, Zhang Y. LINC00908 negatively regulates microRNA-483-5p to increase TSPYL5 expression and inhibit the development of prostate cancer. Cancer Cell Int. 2020 Jan 9;20:10.

52. Sandovici I, Fernandez-Twinn DS, Campbell N, Cooper WN, Sekita Y, Zvetkova I, et al. Overexpression of Igf2-derived Mir483 inhibits Igf1 expression and leads to developmental growth restriction and metabolic dysfunction in mice. Cell Rep. 2024 Sep 24;43(9):114750.

53. Perimbeti S, Jamroze A, Attwood K, Farmer B, Beumer JH, Bies R, et al. Phase Ib trial of enzalutamide (Enza) with venetoclax (Ven) in metastatic castration-resistant prostate cancer (mCRPC). JCO. 2023 Feb 20;41(6_suppl):182–182.

54. Afshar AS, Xu J, Goutsias J. Integrative identification of deregulated miRNA/TF-mediated gene regulatory loops and networks in prostate cancer. PLoS ONE. 2014 Jun 26;9(6):e100806.

55. Oh-Hohenhorst SJ, Lange T. Role of Metastasis-Related microRNAs in Prostate Cancer Progression and Treatment. Cancers (Basel). 2021 Sep 6;13(17).

56. Shalem O, Sanjana NE, Hartenian E, Shi X, Scott DA, Mikkelson T, et al. Genome-scale CRISPR-Cas9 knockout screening in human cells. Science. 2014 Jan 3;343(6166):84–7.

57. Kozomara A, Griffiths-Jones S. miRBase: annotating high confidence microRNAs using deep sequencing data. Nucleic Acids Res. 2014 Jan;42(Database issue):D68–73.

58. Wang T, Birsoy K, Hughes NW, Krupczak KM, Post Y, Wei JJ, et al. Identification and characterization of essential genes in the human genome. Science. 2015 Nov 27;350(6264):1096–101.

59. Morgens DW, Wainberg M, Boyle EA, Ursu O, Araya CL, Tsui CK, et al. Genome-scale measurement of off-target activity using Cas9 toxicity in high-throughput screens. Nat Commun. 2017 May 5;8:15178.

60. Langmead B, Salzberg SL. Fast gapped-read alignment with Bowtie 2. Nat Methods. 2012 Mar 4;9(4):357–9.

61. Tastsoglou S, Alexiou A, Karagkouni D, Skoufos G, Zacharopoulou E, Hatzigeorgiou AG. DIANA-microT 2023: including predicted targets of virally encoded miRNAs. Nucleic Acids Res. 2023 Jul 5;51(W1):W148–53.

62. Perelman A, Wachtel C, Cohen M, Haupt S, Shapiro H, Tzur A. JC-1: alternative excitation wavelengths facilitate mitochondrial membrane potential cytometry. Cell Death Dis. 2012 Nov 22;3(11):e430.

63. Martin M. Cutadapt removes adapter sequences from high-throughput sequencing reads. EMBnet j. 2011 May 2;17(1):10.

64. Li H, Durbin R. Fast and accurate short read alignment with Burrows-Wheeler transform. Bioinformatics. 2009 Jul 15;25(14):1754–60.

65. Pastrello C, Kotlyar M, Abovsky M, Lu R, Jurisica I. PathDIP 5: improving coverage and making enrichment analysis more biologically meaningful. Nucleic Acids Res. 2024 Jan 5;52(D1):D663–71.

66. Subramanian A, Tamayo P, Mootha VK, Mukherjee S, Ebert BL, Gillette MA, et al. Gene set enrichment analysis: a knowledge-based approach for interpreting genome-wide expression profiles. Proc Natl Acad Sci USA. 2005 Oct 25;102(43):15545–50.

67. Mootha VK, Lindgren CM, Eriksson K-F, Subramanian A, Sihag S, Lehar J, et al. PGC-1alpha-responsive genes involved in oxidative phosphorylation are coordinately downregulated in human diabetes. Nat Genet. 2003 Jul;34(3):267–73.

68. Rouillard AD, Gundersen GW, Fernandez NF, Wang Z, Monteiro CD, McDermott MG, et al. The harmonizome: a collection of processed datasets gathered to serve and mine knowledge about genes and proteins. Database (Oxford). 2016 Jul 3;2016.

69. Dreos R, Ambrosini G, Cavin Périer R, Bucher P. EPD and EPDnew, high-quality promoter resources in the next-generation sequencing era. Nucleic Acids Res. 2013 Jan;41(Database issue):D157–64.

70. Raney BJ, Barber GP, Benet-Pagès A, Casper J, Clawson H, Cline MS, et al. The UCSC Genome Browser database: 2024 update. Nucleic Acids Res. 2024 Jan 5;52(D1):D1082– 8.

