## Supplementary figures and images for "A microRNA CRISPR screen reveals miR-483-3p as an apoptotic regulator in prostate cancer"

### Figure S1

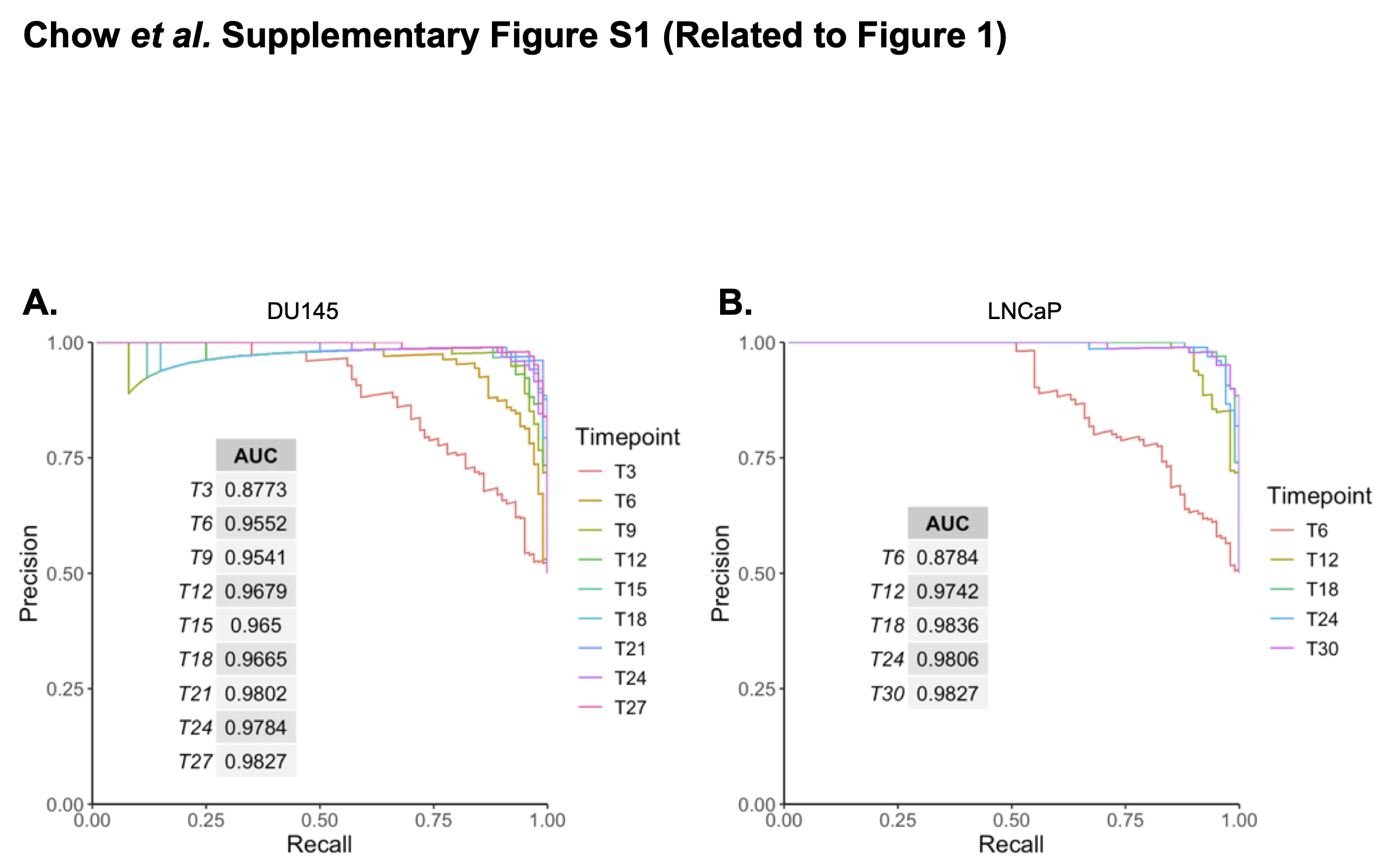

### Figure S2

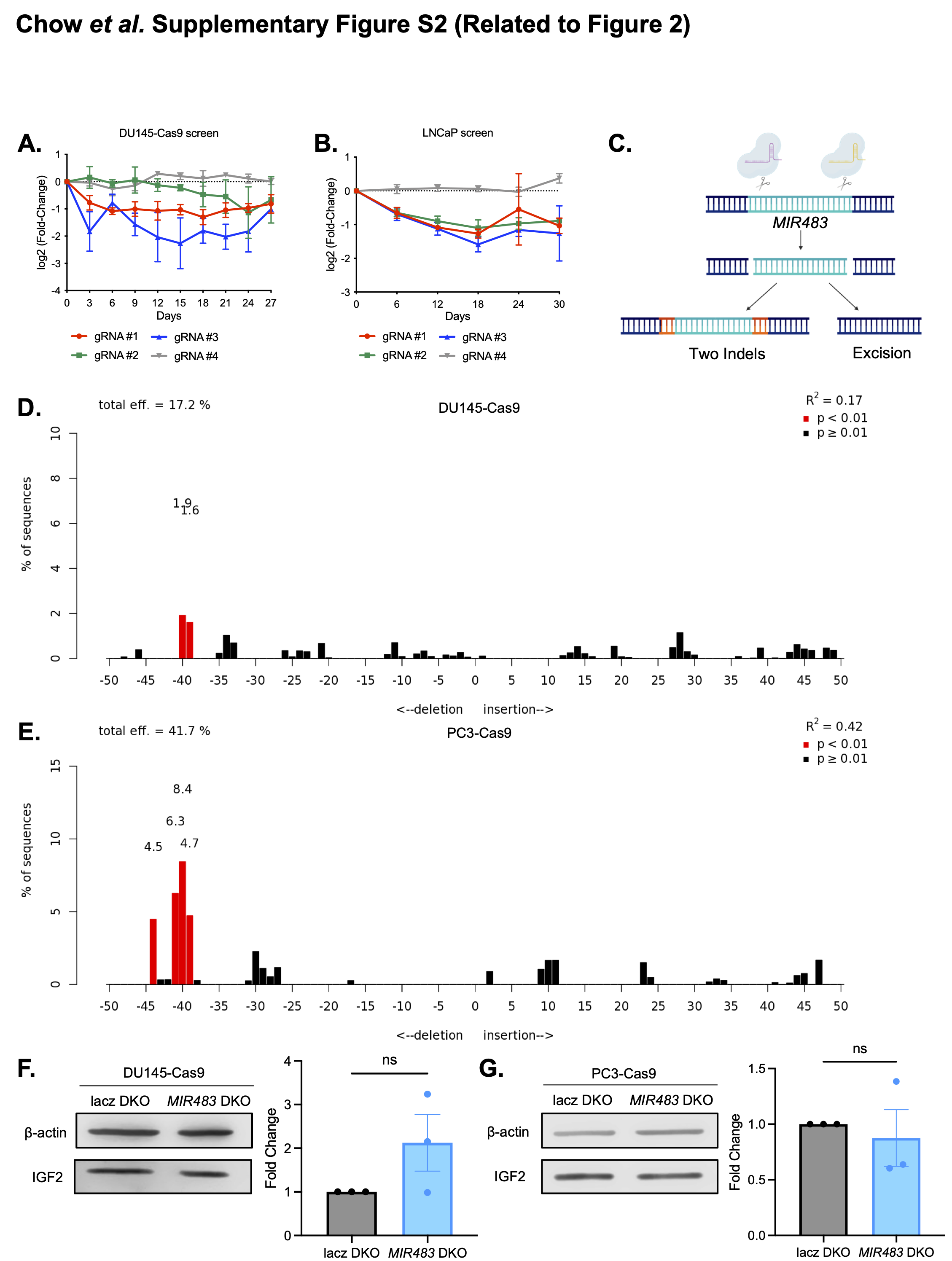

### Figure S3

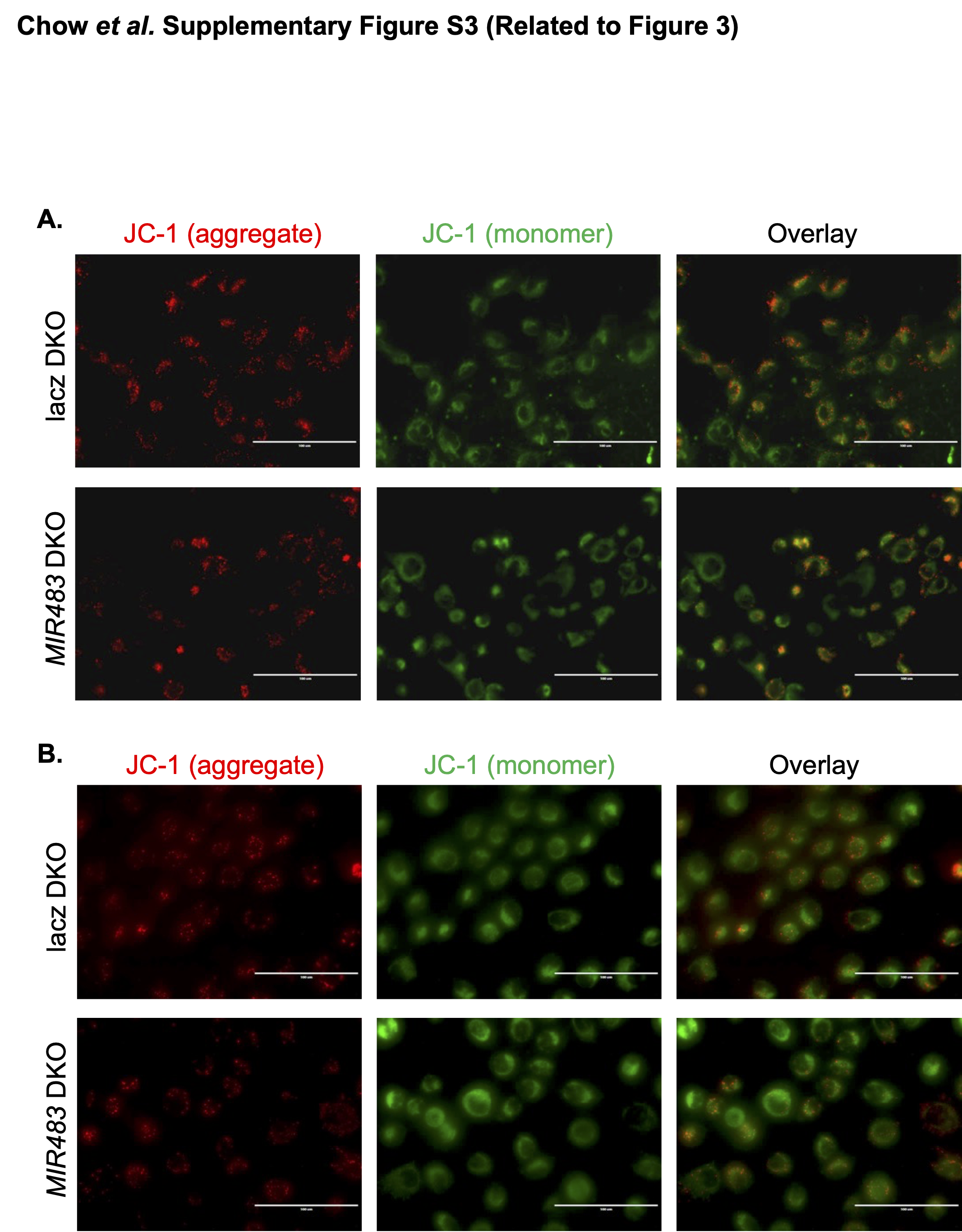

### Figure S4

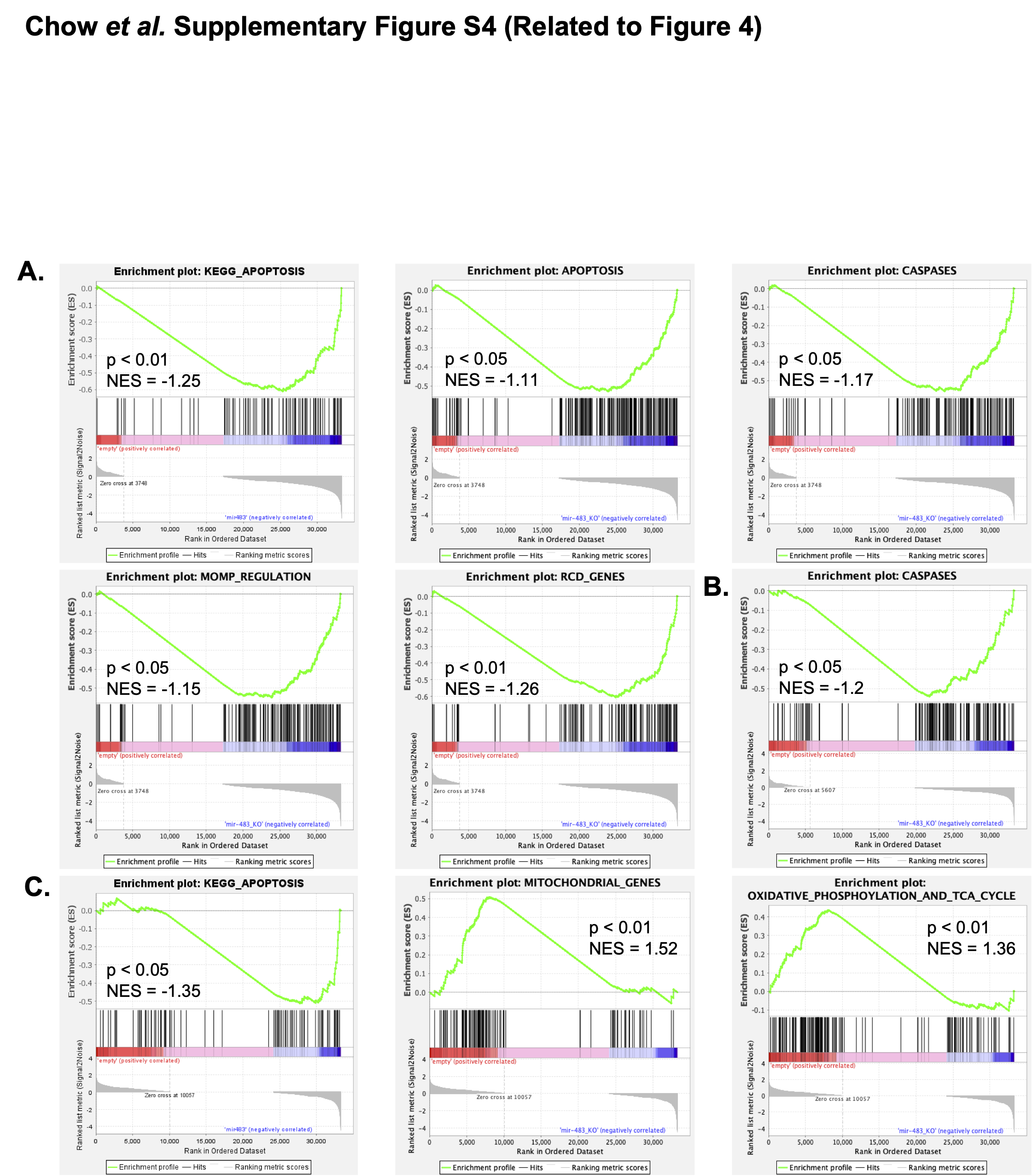

### Figure S5

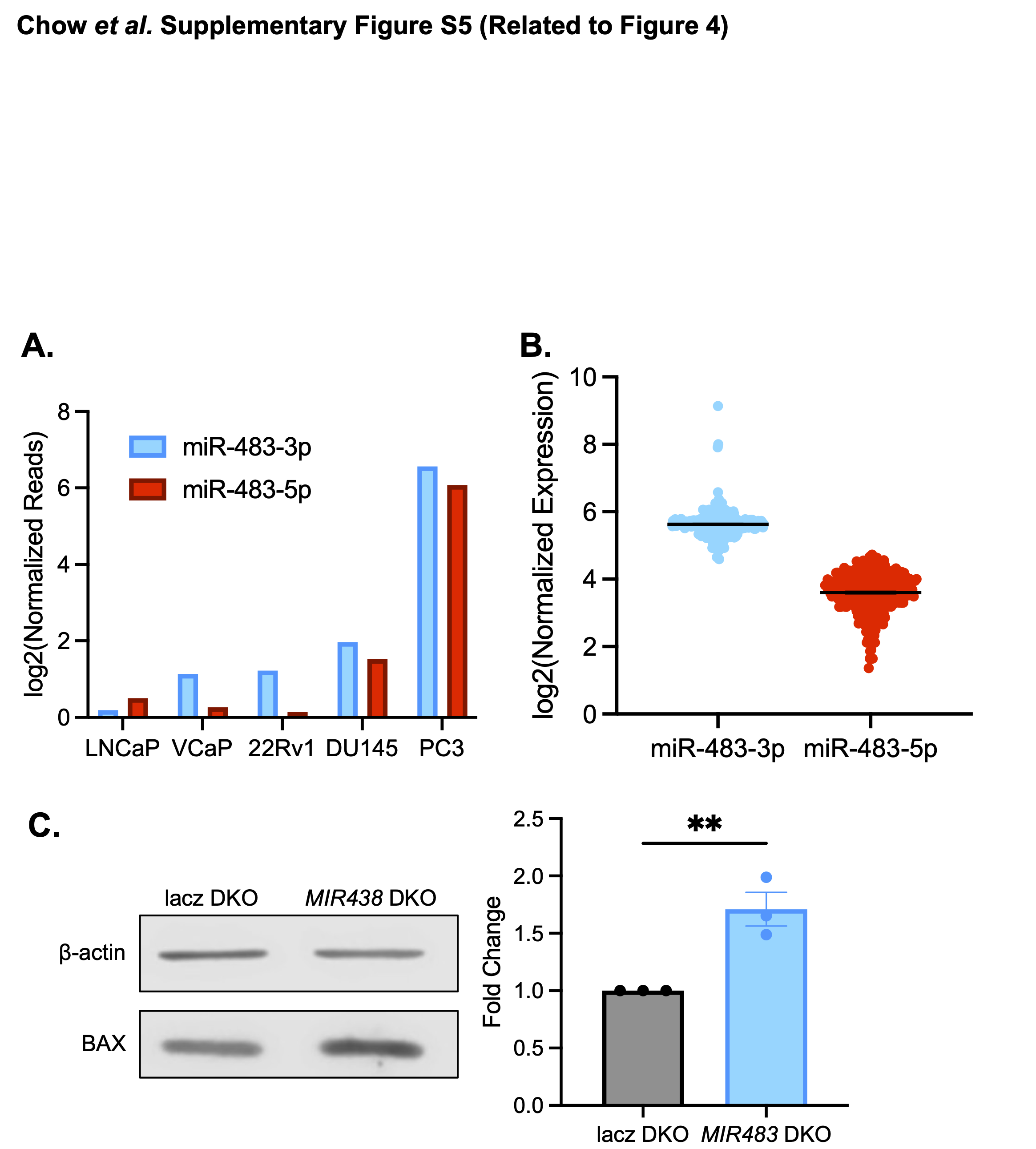

### Figure S6

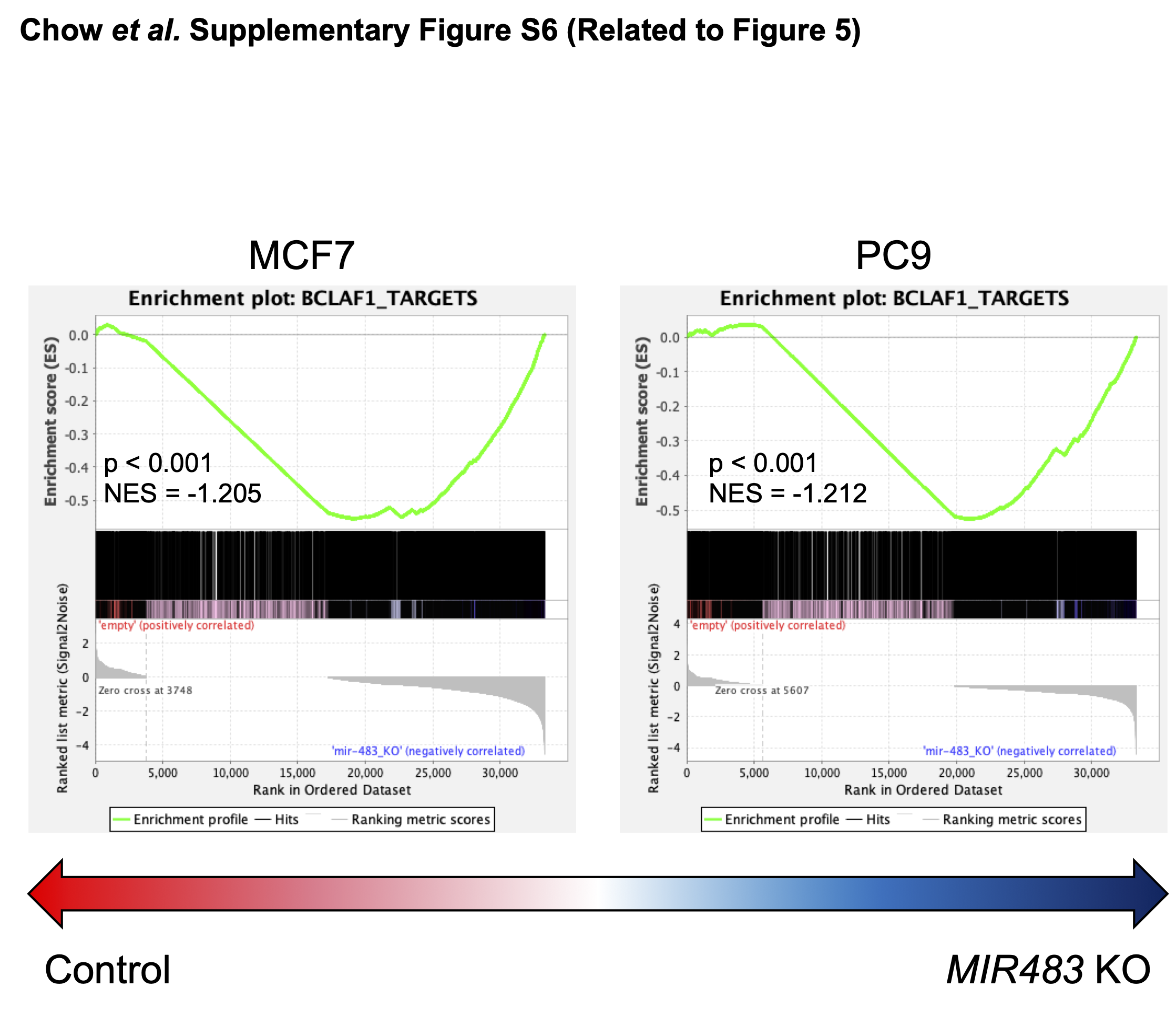
